# Molecular and cellular adaptations in hippocampal parvalbumin neurons mediate behavioral responses to chronic social stress

**DOI:** 10.1101/2021.09.14.459024

**Authors:** Dionnet L. Bhatti, Lucian Medrihan, Michelle X. Chen, Junghee Jin, Kathryn McCabe, Wei Wang, Estefania P. Azevedo, Jose H. Ledo, Yong Kim

## Abstract

Parvalbumin-expressing interneurons (PV neurons) maintain inhibitory control of local circuits implicated in behavioral responses to environmental stressors. However, the roles of molecular and cellular adaptations in PV neurons in stress susceptibility or resilience have not been clearly established. Here, we show behavioral outcomes of chronic social defeat stress (CSDS) are mediated by differential neuronal activity and gene expression in hippocampal PV neurons in mice. Using *in vivo* electrophysiology and chemogenetics, we find increased PV neuronal activity in the ventral dentate gyrus is required and sufficient for behavioral susceptibility to CSDS. PV neuron-selective translational profiling indicates mitochondrial oxidative phosphorylation is the most significantly altered pathway in stress-susceptible versus resilient mice. Among differentially expressed genes associated with stress-susceptibility and resilience, we find Ahnak, an endogenous regulator of L-type calcium channels which are implicated in the regulation of mitochondrial function and gene expression. Notably, Ahnak deletion in PV neurons impedes behavioral susceptibility to CSDS. Altogether, these findings indicate behavioral effects of chronic stress can be controlled by selective modulation of PV neuronal activity or a regulator of L-type calcium signaling in PV neurons.

## INTRODUCTION

Stress can enhance motivational drives that promote evolutionarily favorable behaviors critical for survival (1). However, stressful life events or chronic exposure to unavoidable stress can lead to maladaptive cellular and behavioral responses and contribute to the etiology of neuropsychiatric disease such as major depressive disorder (MDD) (2-6). Many stress-related mental disorders are characterized by altered behavioral and physiological states detrimental to an individual. However, while stress exposure can render some individuals susceptible to developing MDD, others are resilient and often remain healthy (7). Several animal models including ‘learned helplessness’, chronic unpredictable stress, restraint stress and chronic social defeat stress (CSDS) have been used for the studies of stress susceptibility and resilience (8-10). Among them, the CSDS paradigm has been widely used and led to studies that attribute this individual divergence to molecular and cellular adaptations in multiple brain regions (11, 12). Nonetheless, the key molecular and cellular targets responsible for behavioral responses to stress are not fully understood.

Alterations of ventral hippocampus are highly implicated in stress and emotional responses (13, 14). Particularly, multiple neuronal types including granule cells, mossy cells and interneurons in the dentate gyrus (DG) have been studied in regard to depression-like behavior and antidepressant action (15-18). Previously, we found that molecular alterations selectively in parvalbumin (PV)-expressing GABAergic interneurons could modulate depression-like behavior (19, 20). However, whether adaptations in neuronal firing and molecular expression in PV neurons mediate individual differences in stress vulnerability has not been elucidated.

In this study, we find that susceptibility to CSDS is driven by increased PV neuronal activity in the ventral dentate gyrus (PV^vDG^). Furthermore, PV neuron-selective RNA sequencing revealed that CSDS alters multiple molecular pathways regulating mitochondrial function, protein synthesis and synaptogenesis, which could underlie differences in neuronal activity. Among the dysregulated transcripts in hippocampal PV neurons was mRNA for Ahnak. Previously, Ahnak was characterized as an endogenous regulator of L-type voltage-gated calcium channels in PV neurons (20), cardiomyocytes (21) and T cells(22). L-type calcium signaling is highly implicated in mitochondrial function (23, 24), gene expression (25, 26) and neuronal plasticity (27). Importantly, Ahnak expression in the ventral dentate gyrus and in PV neurons is also required for behavioral susceptibility. Together, this study reveals that molecular and cellular adaptations in hippocampal PV neurons incurred by social stress mediate behavioral susceptibility or resilience.

## METHODS AND MATERIALS

### Animals

All experiments involving animals were approved by The Rockefeller University Institutional Animal Care and Use Committee and were in accordance with the National Institutes of Health guidelines. Floxed Ahnak mice were generated and maintained at The Rockefeller University as described previously (20). Floxed Ahnak mice were crossed with PV-Cre mice (stock no: 008069, The Jackson Laboratory) to generate PV neuron-selective Ahnak KO line. Cre-dependent EGFP-L10a mice (stock no: 024750, The Jackson Laboratory, Bar Harbor, ME) were crossed with PV-Cre mice to generate PV neuron-selective EGFP-L10a line for translating ribosomal affinity purification (TRAP). We produced the progeny of floxed Ahnak, PV-Cre, PV-selective EGFP-L10a and PV-selective Ahnak KO lines by *in vitro* fertilization (IVF) and embryo transfer techniques (Transgenic and Reproductive Technology Center, The Rockefeller University) to provide genotype-and age-matched animals. All mice are male and of C57BL/6 background except CD1 aggressors (strain 022, Charles River, Kingston, NY) used for CSDS. Mice were housed 3–5 per cage with a 12:12-h light/dark cycle and *ad lib* access to food and water. *In vivo* recordings were performed with anesthetized mice. All behavioral tests were performed during the light cycle. All behavioral experiments commenced with male mice aged 8-12 weeks old. Transgenic mice were assigned randomly to experimental stress conditions based on their genotype. For AAV-mediated gene delivery experiments, transgenic mice were randomly assigned to experimental virus groups and then later to experimental stress conditions. Experimenters were not blinded to genotype when conducting or analyzing the experiments except during biochemistry experiments, but mice from each group were evenly assigned to equipment and run in parallel during behavioral tests.

### Stereotaxic surgery

All stereotaxic surgeries were performed on an Angle Two Small Animal Stereotaxic Instrument (Leica Biosystems, Buffalo Grove, IL) with a microinjection syringe pump (UMP3 UltraMicroPump, World Precision Instruments, Sarasota, FL). Male mice (7-8 weeks of age) were anesthetized with a mix of ketamine (100 mg/ml) and xylazine (10 mg/ml). AAV5-hSyn-GFP and AAV5-hSyn-Cre-GFP were obtained from UNC Vector Core, and AAV5-hSyn-DIO-mCherry (50459), AAV5-hSyn-DIO-hM4D(Gi)-mCherry (44362) and AAV5-hSyn-DIO-hM3D(Gq)-mCherry (44361) were purchased from Addgene. Viruses (500 μl/side) were injected bilaterally into the vDG (AP -2.7 ML+/-2.0 DV -2.2, mm relative to bregma) with a 2 mL Hamilton Neuros syringe at a speed of 0.1 μl/min. The needle was left for an additional 10 min and then slowly withdrawn. The stereotaxic injections were confirmed by immunohistochemistry and animals with infection outside of vDG or no infection in the vDG were excluded from data analysis. Mice were monitored for 48 h to ensure full recovery from the surgery. Experiments commenced 3 weeks after stereotaxic surgery to allow optimal expression of AAV viruses. Previous studies indicate that a single low dose of ketamine has a long-lasting antidepressant-like effects up to 8 days in rodents (28-30). In this study, we used ketamine as anesthesia for stereotaxic surgeries and AAV-mediated gene delivery. Because we performed experiments 3 weeks after stereotaxic surgery, we believe any potential effect of ketamine had waned down before the commencing the experiments. Importantly, all control groups in Figure 2 and Figure 5A-E underwent the same anesthetic procedure and stereotaxic injection of control AAV, suggesting that the outcome we observed are due to DREADD (designer receptor exclusively activated by designer drugs)-mediated inhibition or activation (Figure 2) or Ahnak KO and not influenced by ketamine.

### Chronic social defeat stress

Chronic social defeat stress (CSDS) was carried out as described previously (31). Retired male breeder CD-1 mice were screened over three consecutive days and aggressors were selected according to the following criteria: (i) the latency to the initial attack was under 60 s and (ii) the screener mouse was attacked for 2 consecutive days. For ten consecutive days, the experimental mice were placed in the home cage of a prescreened CD-1 aggressor for 5-min of physical attack and then separated by a perforated divider for the remaining 24 h until the next defeat 24 h later. Each experimental mouse was exposed to a different aggressor each day. In parallel, stress-naïve control mice were placed in pairs within an identical home cage setup separated by a perforated divider for the duration of the defeat sessions. They were never in physical or sensory contact with CD-1 mice. After 10 days of social defeat, all aggressors and experimental mice were separated and singly housed. The SI test was performed 24□h later.

### Subthreshold social defeat stress

Subthreshold social defeat stress (SSDS) was carried out as described previously (32). Experimental mice were placed in the home cage of a prescreened CD-1 aggressor for 5-min of physical attack. 15 min later, the experimental mice were introduced to another 5-min physical attack by a novel CD-1 aggressor. This was repeated once more for a total of three defeat sessions. The SI test was performed 24□h later.

### Social Interaction Test

Social interaction (SI) test was carried out as described previously (31). The test was composed of two phases, each consisting of 150 seconds, where the experimental mice were allowed to explore an open field (42 cm x 42 cm x 42 cm) with a wire mesh enclosure (10 cm wide x 6.5 cm deep x 42 cm high). In the first phase, the wire mesh enclosure was empty. In the second phase, a novel CD-1 aggressor mouse was placed inside the wire mesh. The amount of time the experimental mice spent in the interaction zone surrounding the wire mesh enclosure was collected and analyzed by the video-tracking apparatus and software EthoVision XT 7 (Noldus Information Technology, Leesburg, VA). SI ratio was calculated by dividing the amount of time the experimental mice spent in the interaction zone in phase two by the time in phase one [SI Ratio = (phase 2 time)/(phase 1 time)]. Susceptible mice were defined by a SI ratio under 1, whereas resilient mice were defined by a SI ratio greater than 1 (31).

### Sucrose Preference Test

Sucrose Preference Test (SPT) was adapted from previous studies (20). During the 1-day habituation period, mice were given a choice of two water bottles. The following day, bottles were replaced with new bottles containing either water or 2% sucrose solution. The consumption of water and sucrose solution was measured at different timepoints (0 and 4 h for acute CNO experiment or 24□h for chronic CNO and genetic knockout experiments). The sucrose preference was represented as percent preference for sucrose ((sucrose consumed/ (water+sucrose consumed)) X 100).

### Drug Administration

For *in vivo* electrophysiology experiments, susceptible mice were selected after the SI test. Ketamine administration (10 mg/kg, i.p., k2753, Sigma-Aldrich) occurred 24 h after the SI test and 1 day before the *in vivo* recording experiment.

For all Gi and Gq DREADD experiments, CNO (3 mg/kg, i.p., C0832, Sigma-Aldrich) (33) was administered. For chronic CNO administration in h4MDi animals during CSDS (Figure 4c), CNO was administered 30 min prior to each defeat for all 10 days of defeat. SI test was performed drug-free. For CNO administration in hM4Di animals after CSDS (Figure 4h), CNO was administered 30 min prior to the SI test and SPT. For CNO administration in hM3Dq animals during SSDS (Figure 4m), CNO was administered 30 min prior to the first subthreshold defeat session. SI-1 test was performed drug-free. For chronic CNO administration in h3MDq animals (Figure 4M), CNO was administered once each day for 10 days beginning the day after the acute administration and SI-1 test. SI-2 test was performed drug-free.

### *In Vivo* Electrophysiology

Animals were anesthetized by injection of urethane (1mg/kg, i.p.). For craniotomy, mice were mounted in a stereotaxic frame (David Kopf Instruments, CA), in which the head of the animal was fixed with a pair of ear bars and a perpendicular tooth bar. Body temperature was continuously monitored by a rectal thermometer and maintained at 33 ± 1 °C by placing the animal on a heating pad. Measurements were obtained from the ventral hippocampus. Stereotaxic coordinates (in mm, anterioposterior [-2.92] measured from bregma; lateral [±2.00] specified from midline; dorsoventral [-2.20] from surface of the brain) were set according to the Franklin and Paxinos Mouse Brain atlas (3rd edition). High density silicon probes with 4 shanks were used (Buzsaki32, NeuroNexus Inc., MI). The shanks were 250 μm apart from each other and bore 8 recording sites each (160 μm diameter for each site; 1-3 MΩ impedance) arranged in a staggered configuration with 20 μm vertical separation. The probes were connected to a RHD 2132 amplifier board with 32 channels (Intan Technologies, CA), mounted on a micromanipulator (Luigs & Neumann, Germany) and were gently inserted in the craniotomy window targeting the granule cell layer of the dentate gyrus in ventral hippocampus (AP –2.9 mm, L 2 mm, DV –2.2 mm). Data were sampled at 20 KHz.

For the spike detection, we employed a fully automated approach with the MountainSort clustering software (publically available at https://github.com/flatironinstitute/mountainlab). Spikes were verified using the MountainView software, available in the same package, and the various parameters were imported in Matlab for the spike sorting procedure (Matlab Signal Processing Toolbox, Mathworks, MA). For the spike sorting procedure, two features of the spike were initially computed: trough-to-peak latency and the bursting behavior. Narrow waveform neurons, likely putative parvalbumin-positive basket cells (34), were classified as neurons with trough-to-peak smaller than 0.4ms; wide waveforms were classified as neurons with half peaks longer than 0.4ms. For the bursting behavior, a bursting index was calculated from the ratio of the frequency distribution of inter-spike intervals at 0-10 ms to that of the frequency distribution of inter-spike intervals at 200-300 ms. Neurons with a bursting index higher than 1.8 were considered excitatory neurons (35). To further separate the excitatory neurons, we took advantage of our own previous experience with intracellular recording with mossy cells and granule cells (36). The total number of neurons detected in all analyzed animals in the study (n=772) went through a first stage classification based on the through-to-peak latency and burst index (BI). Three divisions were classified: putative excitatory cells (black dots, n = 336, BI > 1.8), narrow-waveform (red dots, n = 142, latency ≤ 0.4 ms) and wide-waveform (green dots, n = 294, latency > 0.4 ms) putative interneurons. Based on the bimodal distribution of the afterhyperpolarization (AHP) current, measured from the first derivative of the spike, excitatory neurons were further classified in putative mossy cells (n=133, AHP < 70) and putative granule cells (n=203, AHP > 70). In patch-clamp experiments, the intracellular action potentials of mossy cells show a significantly smaller AHP when compared to the mossy cells (L.M., unpublished data). To analyze the AHP in our experiments, we used the first derivative of the excitatory spikes (dx=dV/ds) in order to avoid the possible errors induced by the variable distance of the respective cells to the recording electrode. This approach also maximizes the differences in AHP induced by the potentially different dynamics between the ion channel conductances in the two neuronal types. We noticed a bimodal distribution of AHP in the excitatory neurons and according to it, we set a threshold of 70 (µV/ms) as a separation between mossy cells (< 70) and granule cells (>70).

### Slice Electrophysiology

4-week-old mice were euthanized with CO_2_. Following decapitation and removal of the brains, transversal slices (400 μm thickness) were cut using a Vibratome 1000 Plus (Leica Microsystems, IL) at 2 °C in a cutting solution containing (in mM): 87 NaCl, 25 NaHCO_3_, 2.5 KCl, 0.5 CaCl_2_, 7 MgCl_2_, 25 glucose, 75 sucrose and saturated with 95% O_2_ and 5% CO_2_. After cutting, the slices were left to recover for 30–45□min at 35□°C and then for 1□h at room temperature in recording solution (aCSF). The aCSF solution contained (in mM): 125 NaCl, 25 NaHCO_3_, 2.5 KCl, 1.25 NaH_2_PO_4_, 2 CaCl_2_, 1 MgCl_2_ and 25 glucose (bubbled with 95% O2 and 5% CO2). Whole-cell patch-clamp recordings were performed with a Multiclamp 700B/Digidata1550A system (Molecular Devices, CA) and an upright Olympus BX51WI microscope (Olympus, Japan). An individual slice was placed in a recording chamber (RC-27L, Warner Instruments, USA) and constantly perfused with oxygenated aCSF at 24 °C (TC-324B, Warner Instruments, USA) at a rate of 1.5–2.0 ml/min. Whole-cell patch-clamp recordings were obtained from PV neurons identified based on their size, shape and position in the subgranular layer using recording pipettes (Glass type 8250, King Precision Glass, Inc., CA) that were pulled in a horizontal pipette puller (Narishige, NY) to a resistance of 3–4 MΩ and filled with an internal solution containing (in mM): 126 K-gluconate, 4 NaCl, 1 MgSO_4_, 0.02 CaCl_2_, 0.1 BAPTA, 15 glucose, 5 HEPES, 3 ATP, 0.1 GTP (pH 7.3). In order to measure the firing of the PV neurons, steps of 100 pA current were injected from a set starting membrane potential of -70 mV. Properties of the single action potentials were measured from the first action potential induced by the steps of injected current. Data were acquired at a sampling frequency of 50 kHz and filtered at 1 kHz and analyzed offline using pClamp10 software (Molecular Devices, CA).

### Translating Ribosome Affinity Purification (TRAP)

TRAP was conducted as previously described (37, 38). Briefly, PV neuron-selective TRAP mice were subjected to CSDS. Hippocampi from PV-TRAP mice were freshly harvested. Hippocampal homogenates of non-defeat, resilient and susceptible mice were used for immunoprecipitation of EGFP-tagged polysomes from PV neurons, and polysome-attached mRNAs were isolated and RNA was further purified using RNeasy Micro Kit (Qiagen, Hilden, Germany). All RNA samples were validated for high quality using Bioanalyzer RNA 6000 Pico Kit (Agilent, San Diego, CA). 1 ng of total RNA was used to generate full length cDNA using Clontech’s SMART-Seq v4 Ultra Low Input RNA Kit. 1 ng of cDNA was then used to prepare libraries using Illumina Nextera XT DNA sample preparation kit. Libraries with unique barcodes were pooled at equal molar ratios and sequenced on Illumina NextSeq 500 sequencer to generate 150 bp single reads, following manufacture’s protocol. Raw data can be found in the Nation Center for Biotechnology Information (NCBI) Gene Expression Omnibbus (GEO) database (GSE184027).

### Bioinformatics analysis

Following sequencing, adapter and low-quality bases were trimmed by fastp (39) from the raw sequencing files in FASTQ format. Cleaned reads were aligned to the Mus musculus assembly 10 reference genome using STAR version 2.7.1a (40). After alignment, the Fragments Per Kilobase of transcript per Million mapped reads (FPKM) for all genes in each sample were calculated with R package edgeR(41). To analyze differential gene expression between samples, DESeq2(42) was used, applying the standard comparison mode between two experimental groups. P values were calculated in DESeq2 adjusted for multiple testing using the Benjamini-Hochberg procedure. Ingenuity Pathway Analysis was used to analyze cellular pathways with differentially expressed genes. The P values for DEGs were then uploaded to Ingenuity Pathway Analysis (Qiagen), which was used to analyze the cellular pathways determined by the differentially expressed genes.

### Quantitative PCR (qPCR)

2.5 ng/2µl of resulting cDNA from TRAP library preparation was used for each qPCR reaction with 10 µl TaqMan Fast Advanced Master Mix (Applied Biosystems), 1 µl of Ahnak Prime Time qPCR primer (Integrated DNA Technologies, Mm.PT.56a.13518996) or 1 µl of Actb Prime Time qPCR primer (IDTDNA, Mm.PT.58.28904620.g), and 7 µl of water. Samples were heated to 50 °C for 2 min, 95 °C for 10s, followed by 40 cycles of 95 °C for 15s, 60 °C for 1 min. Samples were normalized to Actb as a housekeeping gene. mRNA levels were expressed using the 2™ΔΔCt method (43).

### Immunohistochemistry

Animals were deeply anesthetized using CO_2_ and transcardially perfused with PBS, followed by 4% paraformaldehyde (PFA) in PBS. Brains were post-fixed in 4% PFA overnight at 4□°C, and then cryoprotected using 30% sucrose in PBS for at least 24□h, followed by freezing and embedding in Tissue Tek OCT medium (Sakura Finetek USA Inc., CA). A cryostat was used to collect 40-μm-thick coronal sections. All staining between groups used the same master solution mix of blocking buffer and antibodies. Immunohistochemistry was performed side by side between groups. Free-floating sections were washed in PBS and subsequently incubated in blocking buffer (0.5% Triton X-100, 5% normal goat serum, in PBS) for ∼2□h at room temperature. Sections were then incubated overnight (∼16□h) at 4□°C in the primary antibodies diluted in blocking buffer. The primary antibodies were as follows: anti-eGFP (chicken polyclonal, GFP-1020, Aves, 1:1,000), anti-Cre recombinase (mouse monoclonal, MAB3120, Millipore, 1:200), anti-parvalbumin (mouse monoclonal, PV235, Swant, 1:1,000 or guinea pig polyclonal, GP72, Swant, 1:1,000) and anti-mCherry (mouse monoclonal, 3A11, DSHB (UIOWA), 1:500). After incubation, sections were washed three times in PBS and incubated with Alexa-fluor-conjugated secondary antibodies (goat anti-guinea pig, A-11073, Invitrogen, 1:5,000; goat anti-rabbit (31460) and goat anti-mouse (31430), Thermo Fisher Scientific, 1:5,000). After secondary incubation, sections were washed in PBS three times and mounted on glass slides with hard set Vectashield (Vector Labs, CA) for microscopy. Confocal images were obtained on a Zeiss LSM 710 confocal imaging system (Carl Zeiss Microscopy, Thornwood, NY) using a 20×□/0.8□N.A. air or a 100×□/1.4□N.A. oil-immersion objectives (Carl Zeiss Microscopy, Thornwood, NY). Gain, exposure time, and all other related settings were constant throughout each experiment. All image groups were processed in parallel using Fiji.

### Western Blot

Mouse hippocampal tissues were lysed with a lysis buffer (Pierce IP Lysis Buffer, 87788, Thermo Fisher Scientific) supplemented with a protease and phosphatase inhibitor cocktail (78442, Thermo Fisher Scientific). The tissue lysates were homogenized with a Tissue Grinder (10 strokes, 02-911-529, Thermo Fisher Scientific) and centrifuged at 800□X□g for 5□min. Protein levels in the supernatant were measured by the BCA method. The samples were mixed with the standard protein sample buffer and boiled on a hot plate for 2 min, and subjected to SDS-PAGE with 4–20% Novex Tris-Glycine gels (Sigma-Aldrich), followed by protein transfer onto a nitrocellulose membrane. The membranes were Immunoblotted with incubation of anti-Ahnak (rabbit polyclonal, RU2024, 1:5,000) or Anti-Gapdh (mouse monoclonal, MAB374, Millipore, 1:5,000) primary antibody followed by incubation of horseradish peroxidase-linked goat anti-rabbit or anti-mouse secondary antibody (1: 5,000, Thermo Fisher Scientific). Antibody binding was detected using the enhanced chemiluminescence immunoblotting detection system (Perkin Elmer LLC WESTERN LIGHTNING PLUS-ECL, Thermo Fisher Scientific) and Kodak autoradiography film. The bands were quantified by densitometry using NIH Image 1.63 software.

### RNAscope

One day following SI tests, animals were anesthetized and rapidly decapitated. Brains were quickly removed and fresh frozen in dry ice. Sections were cut at 20µm, mounted on slides, and stored at -80°C. Sections were fixed in 4% PFA for 15 min, dehydrated in serial ethanol concentrations (50%, 70%, and 100%) and processed with the RNAscope Multiplex Fluorescent assay (320293, RNAScope, Advanced Cell Diagnostics, Inc., CA). Sections were hybridized with a mixture of selective probes for parvalbumin (Mm-Pvalb, 421931) and Ahnak (Mm-Ahnak, 576971). Sections were then counterstained with DAPI and coverslipped. Confocal images were obtained on a Zeiss LSM 710 confocal imaging system using a 100×□/1.4□N.A. oil-immersion objective (Carl Zeiss Microscopy). Because each image contained only a few cells, multiple stack images were taken of each vDG and treated independently. Z-stacks were taken at 1µm step size. Gain, exposure time, and all other related settings were constant for each quantified image. To generate a projection image for each PV cell, each set of stack projections was z-stacked with maximum intensity using Fiji. Circular ROIs were drawn around all PV-expressing cells in the vDG. These ROIs were then used for puncta and cell count analysis using Fiji. Total n=73-102 PV cells in vDG were analyzed per group. 2-3 sections per mouse and total 3-4 mice per group were analyzed.

### Statistics

All data are expressed as means ± SEM except violin plots and pie charts. Sample sizes for biochemistry, electrophysiology and CSDS were determined based on our empirical data accumulated in the laboratory and previous studies using CSDS (10, 44, 45). Sample sizes and statistical methods are provided in each figure legend. Statistical analysis was performed using the two-tailed unpaired Student’s t-test, one-way ANOVA, two-way ANOVA or repeated-measures two-way ANOVA (see Summary of statistical analysis in Supplementary Table 1). ANOVAs of paired Gaussian distributions were followed with Bonferroni’s post-hoc test. The D’Agostino-Pearson omnibus normality test was used to determine normality. For non-Gaussian distributions of multiple groups, the Kruskal-Wallis test followed by Dunn’s multiple comparisons post-hoc test was used. GraphPad Prism 6 or 8 was used for statistical analysis and graphical preparations. p < 0.05 was considered the cutoff for statistical significance. Statistical significance is shown as *p<0.05, **p<0.01, ***p<0.001, ****p<0.0001, otherwise indicated in figures or figure legends.

## RESULTS

### PV neuron activity in vDG is associated with behavioral responses to CSDS

The vDG is involved in mood control, stress responses, and antidepressant actions (18, 46, 47). To assess the role of vDG neurons in stress-induced behavioral outcomes, we recorded vDG neuronal activity in mice that underwent CSDS (Figure 1A). The CSDS paradigm uses ethologically relevant stressors including physical defeat by and sensory contact with aggressor mice for 10 days to evoke social avoidance and anhedonic-like behavior, as measured by social interaction (SI) with a novel aggressor mouse and sucrose consumption (10, 31). The SI test produces an SI ratio of the time spent in the interaction zone in the presence or the absence of a novel aggressor to identify susceptible (SI ratio below 1) and resilient (SI ratio equal to and greater than 1) groups (Figure 1B, C, Supplemental Figure S1). Non-defeated control mice remained housed in pairs without social defeat or contact with aggressors and display SI ratios similar to resilient mice.

**Figure 1.**
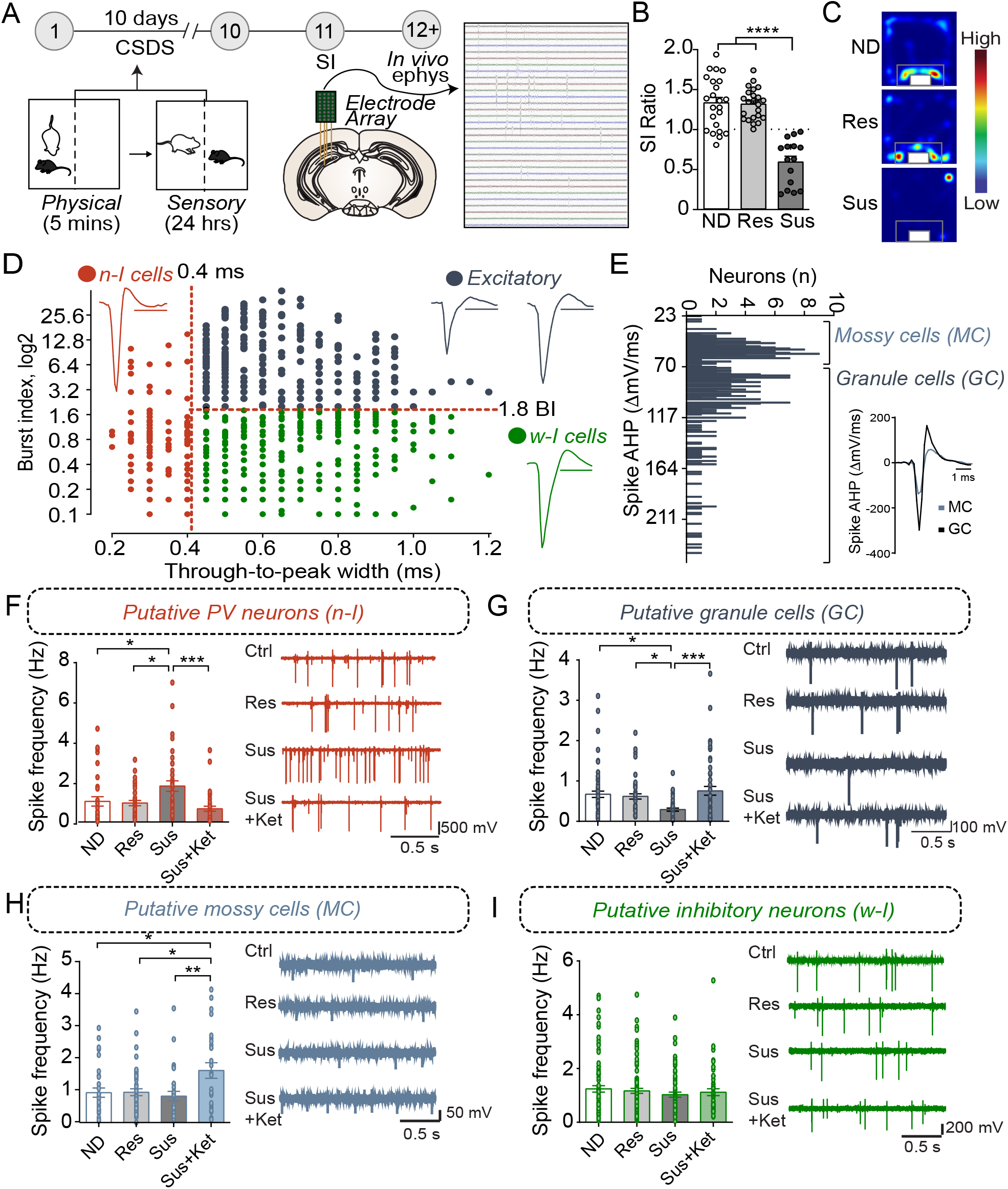
PV^vDG^ neuron activity is altered by CSDS and associated with behavioral outcome. **(A)** Diagram and timeline of CSDS and intra-vDG electrophysiological recordings. **(B)** Representative animals depicting separation of animals into non-defeated control (ND), resilient (Res, SI ratio ≥ 1), and susceptible (Sus, SI ratio < 1) groups (one-way ANOVA, Non-defeated (n=23), Res (n=23) and Sus (n=15)). **(C)** Example heatmaps displaying time spent in the SI arena with caged aggressor. **(D)** All neurons detected (n=772) went through a first stage classification based on the through-to-peak latency and burst index (BI) to distinguish putative excitatory cells (black dots, n = 336, BI > 1.8), narrow-waveform putative PV neurons (red dots, n = 142, latency ≤ 0.4 ms) and wide-waveform putative interneurons (green dots, n = 294, latency > 0.4 ms). Scale bars for the representative waveforms represent 1 ms. **(E)** Excitatory neurons were further classified in putative mossy cells (n=133, AHP < 70) and putative granule cells (n=203, AHP > 70) based on the bimodal distribution of the AHP current, measured from the first derivative of the spike. **(F-I)** Histograms show frequency (mean ± SEM) of the putative neuronal types analyzed in the respective experimental groups. Each black dot represents a neuron. (**F**) Susceptibility is associated with a decrease in PV neuron spike frequency that is reversed by Ketamine (n = 31/7 (neurons/mice) ND mice; 36/7 resilient mice, 37/5 susceptible mice, and 38/3 for susceptible mice treated with ketamine). (**G**) Susceptibility is associated with decreases in GC spike frequency that is reversed by Ketamine (n = 61/7 (neurons/mice) ND mice, 47/7 resilient mice, 41/5 susceptible mice, and 54/3 susceptible mice treated with ketamine). (**H**) Ketamine increases MC spike frequency (n = 31/7 (neurons/mice) ND mice, 49/7 for the resilient mice, 27/5 for the susceptible mice, and 26/3 susceptible mice treated with ketamine. (**I**) CSDS or Ketamine did not alter spike frequency in other inhibitory neurons (n = 85/7 (neurons/mice) ND mice, 97/7 resilient mice, 62/5 susceptible mice and 50/3 susceptible mice treated with ketamine). \**p* < 0.05, \*\**p* < 0.01, \*\*\**p* < 0.001, one-way ANOVA with Bonferroni’s *post hoc* comparison.

To examine whether CSDS elicited adaptations in basal neuronal activity within the vDG, we recorded vDG neurons using *in vivo* high-density silicon probe recordings in anesthetized animals. Harnessing known characteristics of firing dynamics, we classified and separated discrete neuron classes. For example, neurons were classified on the basis of the width (through-to-peak latency) of the waveform of their spikes (Figure 1D. see Methods and Materials section) into ‘narrow’ and ‘wide’ waveform neurons to separate putative fast-spiking PV-positive neurons from other cell classes in the hippocampus (34). Additionally, the analysis of bursting, a marker of internal firing dynamics that distinguishes wide waveform putative excitatory from inhibitory neurons in the vDG, further separated the neurons into wide-inhibitory neurons and excitatory neurons (35). Furthermore, mossy cells have a small AHP current following an action potential (48) compared to granule cells (see Methods) and, based on the bimodal distribution of the amplitude of the AHP current, we further separated the excitatory neurons in putative mossy cells and putative granule cells (Figure 1E).

Our results showed that, in susceptible mice, the firing frequency of putative PV neurons was significantly increased in susceptible mice compared to non-defeated or resilient mice (Figure 1F). In contrast, the firing frequency of putative granule cells was significantly decreased in susceptible mice compared to the frequency in non-defeated or resilient mice (Figure 1G). No difference, however, was found in the firing frequency of mossy cells between non-defeated, susceptible, or resilient groups (Figure 1H). To examine whether CSDS-induced adaptations in neuron activity could be reversed by known fast-acting antidepressants (49), we administered ketamine in susceptible mice and recorded neuronal activity. Ketamine ameliorated the increase in firing frequency of putative PV neurons and restored them to comparable levels to non-defeated or resilient mice (Figure 1F). In addition, ketamine treatment increased firing rates in putative granule and mossy cells (Figure 1G, H). However, the firing frequency of other inhibitory neurons (CCK, SOM etc.) remained unchanged in resilient and susceptible mice with or without ketamine treatment as compared to control mice (Figure 1I). Together, these data suggest that CSDS-induced neuron activity changes of PV^vDG^ and granule cells (GC^vDG^) may play a critical role in divergent behavioral responses to stress. The opposing directionality in CSDS-induced changes in PV neurons and GCs likely arise from the perisomatic inhibition onto large GC populations by PV neurons (50, 51). Our previous studies suggested that inhibition of calcium signaling or neuronal activity selectively in PV neurons induces an antidepressant-like behavioral phenotype (19, 20), and this notion is also supported by the reduction of PV^vDG^ neuron activity by ketamine (Figure 1F). We thus sought to determine whether inhibition or activation of PV^vDG^ was capable of conferring or suppressing resilience to CSDS, respectively.

### PV activity in vDG mediates behavioral responsivity to CSDS

To evaluate whether PV^vDG^ neuronal activity mediates behavioral responses to CSDS, we employed the use of DREADDs (designer receptor exclusively activated by designer drugs) (52) to manipulate PV^vDG^ neuronal activity *in vivo*. The vDG of PV-Cre mice were bilaterally injected with Cre-dependent hM4Di-mCherry or empty vector mCherry (Figure 2A) to selectively express hM4Di-mCherry in PV^vDG^ (Figure 2B). First, we assessed whether repeated chemogenetic inhibition of PV neurons concurrent with CSDS would affect behavioral divergence. Mice expressing hM4Di or mCherry in PV^vDG^ underwent CSDS, during which the DREADD agonist clozapine N-oxide (CNO) was administered 30 minutes prior to each defeat session to suppress neuronal activity during and after each social stress encounter (Figure 2C). Interestingly, repeated inhibition of PV^vDG^ neuronal activity promoted resilience evident from an increased SI ratio (Figure 2D, E, Supplementary Figure 2A). Additionally, repeated CNO-treatment resulted in mitigated stress-induced anhedonic-like behavior in the sucrose preference test (SPT) (10) in mice expressing hM4Di in PV^vDG^ compared to mCherry (Figure 2F).

**Figure 2.**
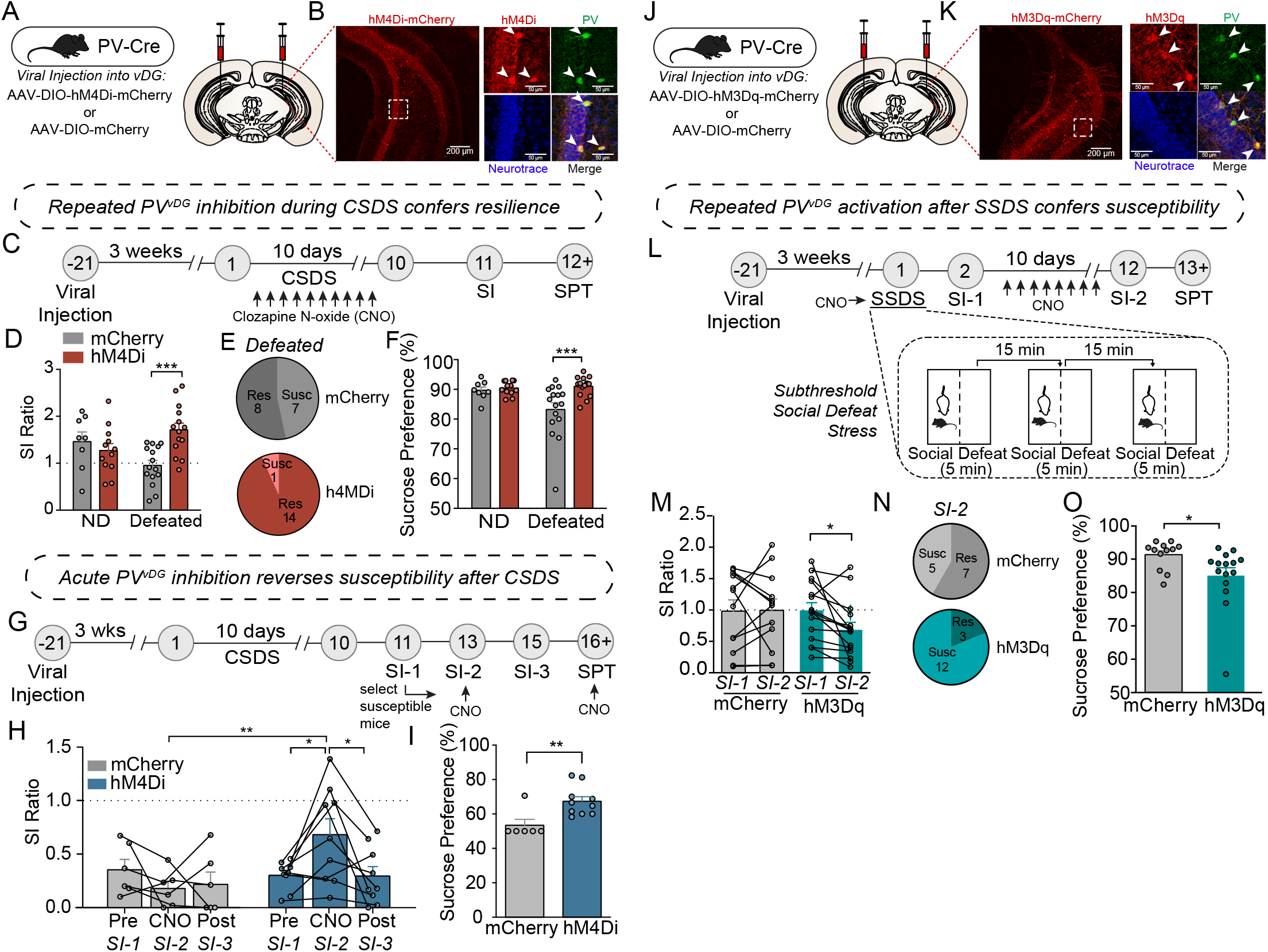
PV neuron activity in the vDG mediates resilience or susceptibility to CSDS. **(A)** Viral injection strategy for expression of hM4Di-mCherry or mCherry in vDG of PV-Cre mice. **(B)** Representative images showing hM4Di-mCherry expression in the vDG (left image; scale bar, 200 μm) and high magnification images of a dotted rectangular region showing selective expression of hM4Di-mCherry in PV neurons (right images; scale bar, 50 μm). **(C)** Timeline of repeated CNO treatments during CSDS and behavioral experiments. **(D)** Repeated CNO-treatment resulted in increased SI ratio (two-way ANOVA, Non-defeat (ND): mCherry (n=8) and hM4Di (n=12), Defeated: mCherry (n=15) and hM4Di (n=14)). **(E)** Pie charts showing the number of resilient and susceptible animals after repeated injections of CNO during CSDS. **(F)** Chronic-CNO treatment during CSDS resulted in hM4Di mice displaying increased sucrose compared to mCherry controls (two-way ANOVA, Non-defeat: mCherry (n=8) and hM4Di (n=12), Defeated: mCherry (n=16) and hM4Di (n=16)). **(G)** Timeline of CSDS, acute CNO treatments and behavioral experiments. **(H)** Acute CNO-treatment resulted in increased SI ratio only in hM4Di but not in mCherry control group during SI-2 test (two-way repeated measures ANOVA, mCherry (n=6) and hM4Di (n=9)). **(I)** After acute-CNO treatment, hM4Di group displayed increased sucrose preference in SPT compared to mCherry control group (unpaired two-tailed t-test, mCherry (n=6) and hM4Di (n=10)). **(J)** Viral injection strategy for expression of hM3Dq-mCherry or mCherry in vDG of PV-Cre mice. **(K)** Representative images showing hM3Dq-mCherry expression in the vDG (left image; scale bar, 200 μm) and high magnification images of a dotted rectangular region showing selective expression of hM3Dq-mCherry in PV neurons (right images; scale bar, 50 μm). **(L)** Timeline of SSDS, CNO treatments and behavioral experiments, and diagram of SSDS. **(M)** Comparison of SI ratio obtained from SI-1 (tested after an acute injection of CNO immediately followed by SSDS) versus SI-2 (tested after repeated daily injections of CNO for 10 days). After chronic injections of CNO, hM3Dq mice displayed increased social avoidance compared to mCherry controls (two-way repeated measures ANOVA, mCherry (n=12) and hM3Dq (n=15)). **(N)** Pie charts showing the number of resilient and susceptible animals resulting from repeated injections of CNO after SSDS. **(O)** After repeated injections of CNO, hM3Dq mice displayed decreased sucrose preference compared to mCherry controls in SPT (unpaired two-tailed *t*-test, mCherry (n=12) and hM3Dq (n=15)). Data are expressed as mean ± SEM, and individual data points are depicted. *Post-hoc* Bonferroni’s multiple comparisons were used for ANOVA. \**p*<0.05, \*\**p*<0.01, \*\*\**p*<0.001.

Given that stress-susceptible mice administered with ketamine displayed PV^vDG^ firing rates similar to non-defeated and stress-resilient mice (Figure 1F), we questioned whether acute inhibition of PV^vDG^ might be sufficient to reverse social avoidance and anhedonic-like behavior in stress-susceptible mice. To address this, mice were exposed to CSDS to generate susceptible mice expressing either hM4Di or empty vector mCherry as a control. 48 hours after the initial SI test (SI-1), only susceptible mice (SI ratio<1.0) were administered with CNO 30 minutes before a subsequent SI-test (SI-2) (Figure 2G). Remarkably, acute CNO administration was capable of attenuating susceptibility as evident by an increased SI ratio (Figure 2H) and time spent in the interaction zone with the aggressor (Supplementary Figure 2B). This amelioration of social avoidance returned to SI scores similar to those in SI-1 when tested another 48-hours later (SI-3) before which CNO was not administered (Figure 2H). To test whether acute inhibition of PV^vDG^ can reverse stress-induced anhedonic-like behavior, we administered CNO during SPT. Indeed, acute administration of CNO increased sucrose preference in mice expressing hM4Di compared to mCherry control mice (Figure 2I).

We next evaluated whether chemogenetic activation of PV^vDG^ would be sufficient to produce stress susceptibility. We bilaterally injected Cre-dependent AAV expressing hM3Dq-mCherry or empty vector mCherry into the vDG of PV-Cre mice (Figure 2J) to selectively express hM3Dq-mCherry in PV^vDG^ (Figure 2K). We adopted a subthreshold social defeat stress (SSDS) paradigm that uses 3 social stress sessions in 1 day (32) and tested the effect of acute activation of PV^vDG^ by CNO during SSDS or repeated activation after SSDS on behavior (Figure 2L). While acute CNO treatment during SSDS did not cause any significant difference between hM3Dq mice and mCherry control mice (Supplementary Figure 2C, D), subsequent repeated daily activation of PV^vDG^ neurons after SSDS resulted in behavioral susceptibility indicated by a decreased SI ratio (Figure 2M, N, Supplementary Figure 2E) and reduced sucrose preference (Figure 2O). Repeated CNO administration in hM3Dq mice naïve to previous stressors, however, failed to elicit this decrease (Supplementary Figure 2F-H). These data suggest that while repeated or acute inhibition is sufficient to drive resilience to social stress, activation of these neurons must occur repeatedly over the course of many days to drive stress-susceptibility, potentially indicative of what occurs during CSDS.

### Molecular adaptations in hippocampal PV neurons are associated with divergent behavioral consequences after CSDS

To determine whether molecular adaptations in hippocampal PV neurons were associated with resilience or susceptibility to CSDS, we generated PV neuron-specific expression profiles using the TRAP/RNA-seq approach(53). We used a Cre recombinase-dependent TRAP line crossed with PV-Cre line to express the ribosomal protein L10a fused to enhanced green fluorescent protein (EGFP) selectively in PV neurons (Figure 3A). After CSDS, mouse hippocampal homogenates of non-defeated, resilient, and susceptible mice were used for immunoprecipitation of EGFP-tagged polysomes from PV neurons, and polysome-attached mRNAs were isolated and sequenced (Figure 3B). Importantly, PV-neuron specific TRAP revealed high selectivity, evident by high enrichment of PV mRNA, but not markers of other neuronal types, neuroglia and cell types in blood vessels, suggesting tight selectivity (Figure 3C).

**Figure 3.**
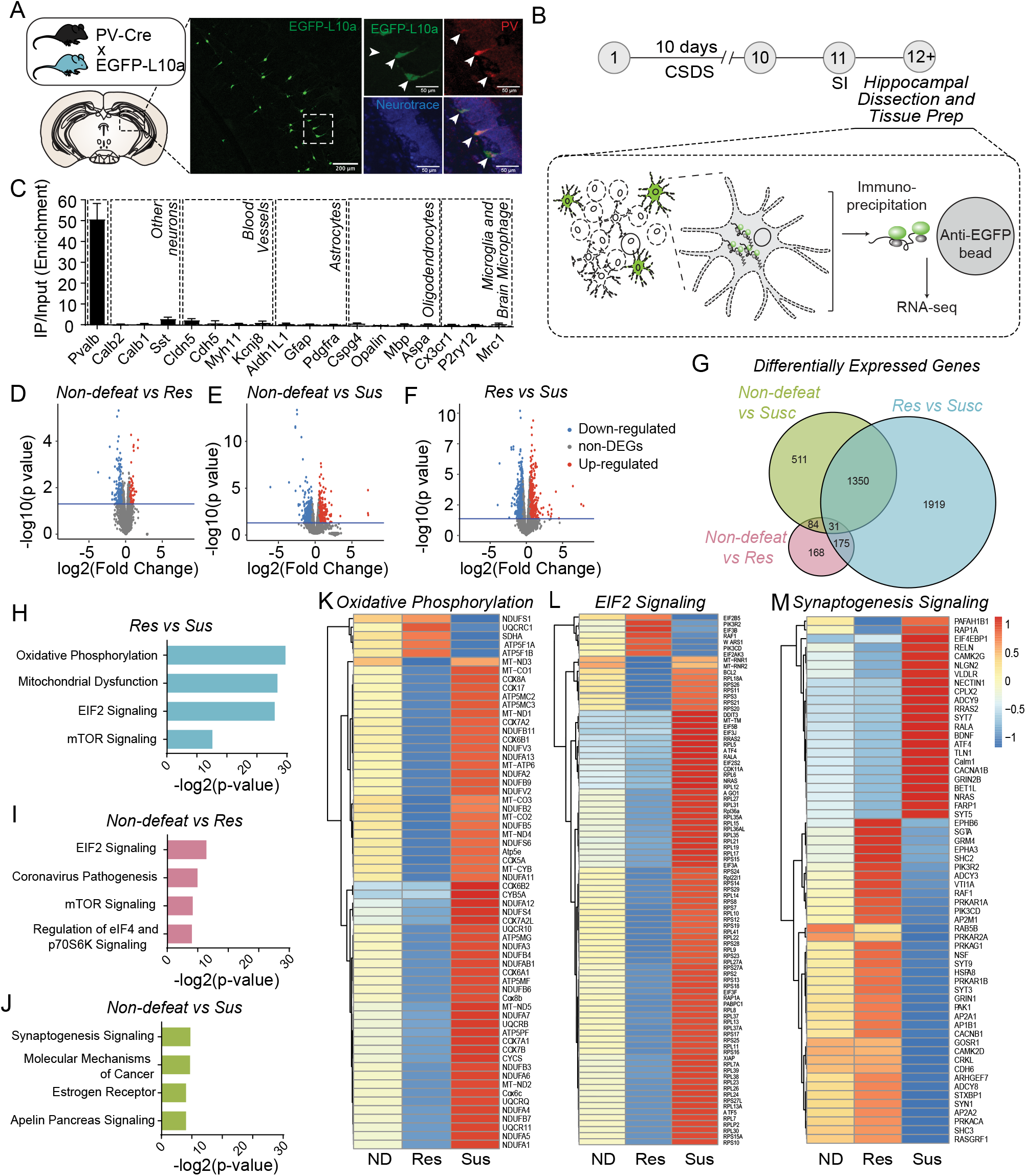
Molecular adaptations in hippocampal PV neurons are associated with divergent behavioral consequences after CSDS. **(A)** Diagram of transgenic PV-specific EGFP-L10a line for TRAP. Representative images showing EGFP-L10a expression in the vDG (left image; scale bar, 200 μm) and high magnification images of a dotted rectangular region showing selective expression of EGFP-L10a in PV neurons (right images; scale bar, 50 μm). **(B)** Timeline of CSDS and subsequent hippocampal dissection, TRAP and RNAseq. **(C)** Enrichment of marker genes in TRAP/RNAseq data. **(D-F)** Volcano plots of DEGs in Non-defeated (ND) vs Res (**D**), ND vs Sus (**E**) and Res vs Sus (**F**) comparisons. **(G)** Venn diagram for DEGs (cutoff, p< 0.05). **(H-J)** Top 4 pathways significantly altered in Res vs Sus **(H)**, ND vs Res (**I**), ND vs Sus (**J**) and comparisons. **(K-M)** Heat maps of DEGs in the top pathway in each comparison. Mean value of gene expression was used (ND (n=4), Res (n=4) and Sus (n=3)).

RNA-seq revealed 458 differentially expressed genes (DEGs) between non-defeated and resilient mice, 1976 DEGs in non-defeated and susceptible mice, and 3475 DEGs in resilient and susceptible mice (Figure 3D-G). Pathway-analysis with DEGs was performed (Figures 3H-J), and between resilient and susceptible mice or non-defeated, genes associated with oxidative phosphorylation (Figure 3K), mitochondrial dysfunction (Supplementary Figure 3B), EF2 signaling (Figure 3L) or mTOR signaling (Supplementary Figure 3B) were differentially expressed. In contrast, changes in expression of genes associated with synaptogenesis were most divergent between non-defeated and susceptible mice (Figure 3J, M). These results suggest that alterations of mitochondrial, protein synthesis, cell metabolism and synaptic pathways in hippocampal PV neurons may drive adaptive changes in PV^vDG^ neuronal activity and behavioral responses to CSDS.

### Chronic stress alters expression of Ahnak in the hippocampus

Our previous study identified Ahnak as a regulator of ‘depressive’-like behavior, potentially in part due to its role in trafficking L-type VGCCs to the surface, positioning Ahnak to regulate neuronal activity responses (20). Remarkably, we find that susceptible mice have increased Ahnak mRNA in our TRAP/RNAseq dataset (p=0.0399, Supplementary Table 2). To confirm whether behavioral responses to CSDS are associated with changes in Ahnak expression, we subjected mice to CSDS followed by SI-test and brain tissue collection (Figure 4A). First, we performed immunoblotting for Ahnak protein in the whole hippocampus. Ahnak protein levels were reduced in the hippocampus of resilient mice but increased in the hippocampus of susceptible mice, compared to non-defeated control mice (Figure 4B-C). Hippocampal Ahnak levels were also inversely correlated with SI ratio (Figure 4D) and time spent in the interaction zone containing an aggressor mouse (Supplementary Figure 4A). These data indicate that CSDS induces alterations of hippocampal expression of Ahnak in opposing directions, depending on the behavioral response of individual mice.

**Figure 4.**
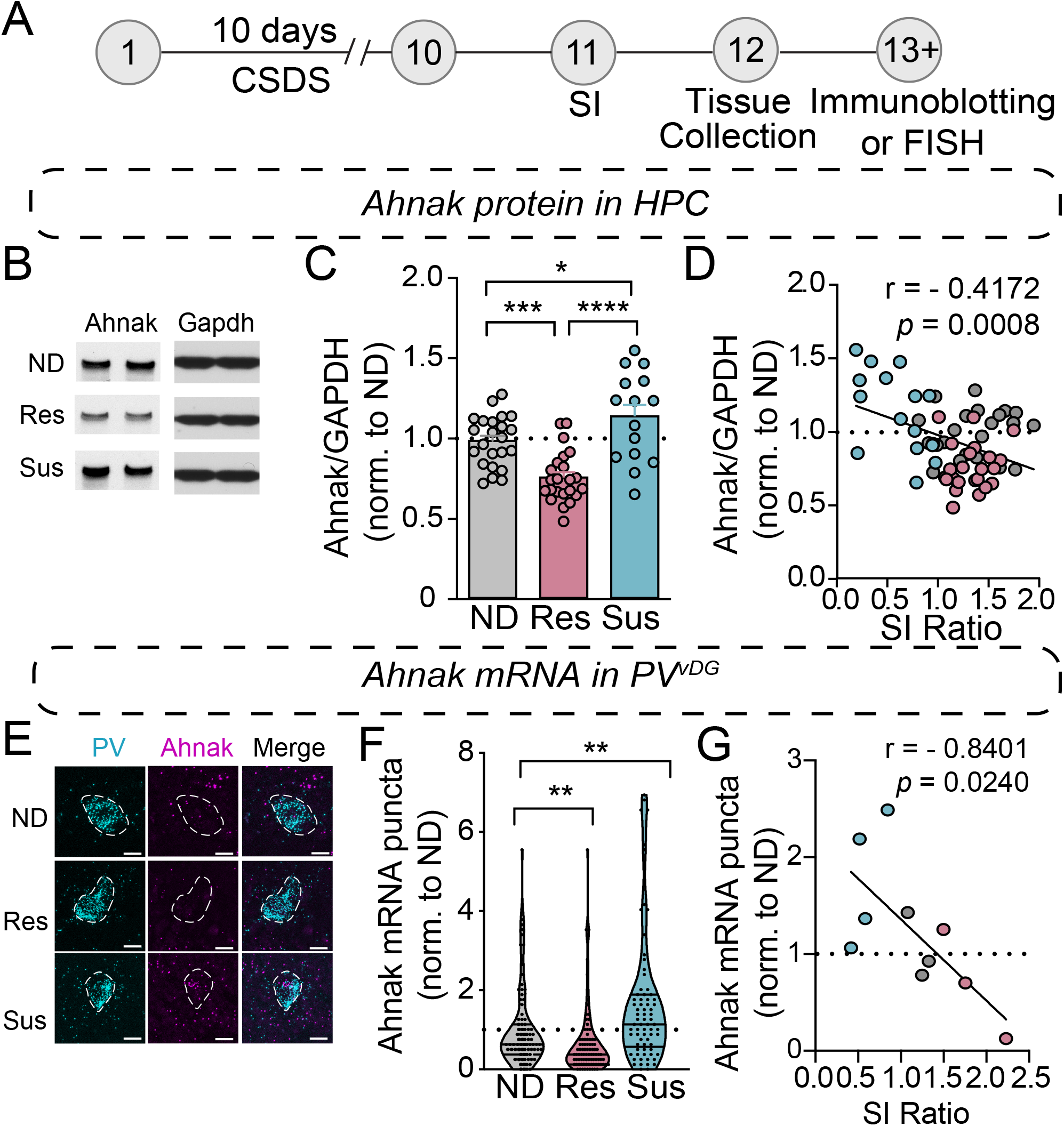
Chronic social stress alters Ahnak expression in the hippocampus. **(A)** Diagram and timeline of CSDS and tissue collection. **(B-D)** Immunoblotting of Ahnak protein using hippocampal lysates. (**B**) Representative images of immunoblotting of Ahnak and Gapdh as a control. (**C**) Quantification of immunoblots (one-way ANOVA, Non-defeated (n=23 mice), Res (n=23 mice) and Sus (n=15 mice)). (**D**) Hippocampal Ahnak is inversely correlated with SI ratio (pearson *r* : *r* = -0.4172, *p*=0.0008, n=61 mice) **(E)** Representative images PV and Ahnak mRNA expression using RNAScope. Scale bar, 5 µm. **(F)** Ahnak mRNA expression in vDG PV neurons from Non-defeat, Res and Sus groups (Kruskal-Wallis test with *post hoc* Dunn’s multiple comparisons: Non-defeated (n=102 cells), Res (n=94 cells) and Sus (n=73 cells). **(G)** Ahnak mRNA expression in vDG PV neurons is inversely correlated with SI ratio (pearson *r* : *r* = -0.8401, *p*=0.0240, n=10 mice). All data were normalized to ND group. \**p*<0.05, \*\**p*<0.01, \*\*\**p*<0.001, \*\*\*\**p*<0.0001.

Our previous study showed that PV neuron-selective Ahnak KO mice displayed antidepressant-like behavior (20). However, homogenates used for immunoblotting include protein of all cell types and subregions within the hippocampus. To evaluate whether CSDS induces alterations of Ahnak expression within PV neurons, we turned to evaluating Ahnak mRNA within PV^vDG^ in non-defeated, stress-resilient and -susceptible mouse brains using RNAscope, a fluorescent RNA *in situ* hybridization assay (54). We used a selective probe for Ahnak mRNA together with a selective probe for PV mRNA in order to indicate the location of PV neurons and quantify Ahnak expression in PV^vDG^ (Figure 4E). The number of puncta of Ahnak per PV^vDG^ was significantly increased in susceptible mice as compared to the number in non-defeated controls or resilient mice, while the resilient mice displayed a significant decrease in Ahnak expression compared to non-defeated control mice (Figure 4F). Additionally, Ahnak mRNA in PV^vDG^ is inversely correlated with SI Ratio and interaction zone containing an aggressor mouse (Figure 4G and Supplemental Figure 4B). This result is consistent with a bidirectional change of Ahnak protein in the hippocampal tissues of susceptible versus resilient mice compared to non-defeated control mice. These results altogether indicate that hippocampal Ahnak expression in PV^vDG^ neurons is altered by CSDS, particularly upregulated in stress-susceptible mice.

### Ahnak deletion in vDG or PV neurons confers behavioral resilience to CSDS

To investigate whether Ahnak is required for CSDS-induced behavioral susceptibility, our approach was two-fold: to assess the behavioral consequence of Ahnak deletion 1) in the vDG and 2) selectively in PV neurons. To delete Ahnak in a region-selective manner, we generated Cre-loxP-mediated ventral dentate gyrus (vDG)-specific Ahnak knockout (KO) mice. We used floxed Ahnak mice (20) injected with adeno-associated virus (AAV) expressing Cre recombinase fused to GFP, or empty vector GFP as a control, under a human synapsin promoter into the ventral dentate gyrus (vDG) to selectively delete Ahnak in vDG neurons (cKO^vDG^) (Figure 5A). Immunohistochemical staining of GFP, Cre recombinase and Ahnak reveal selective KO of Ahnak in neurons infected by AAV-Cre-GFP compared to AAV-GFP (Figure 5B). In response to CSDS, Ahnak cKO^vDG^ resulted in increases of SI ratio (Figure 5C) and time interacting with an aggressor compared to control mice (Supplementary Figure 5A), generating a greater portion of resilient mice (Figure 5D). Additionally, defeated Ahnak cKO^vDG^ mice display mitigated anhedonic-like behavior as measured by increased sucrose consumption compared to defeated control mice (Figure 5E). However, non-defeated Ahnak cKO^vDG^ mice and control mice display comparable behaviors in SI test (Figure 5C and Supplementary Figure 5A) and SPT (Figure 5E).

**Figure 5.**
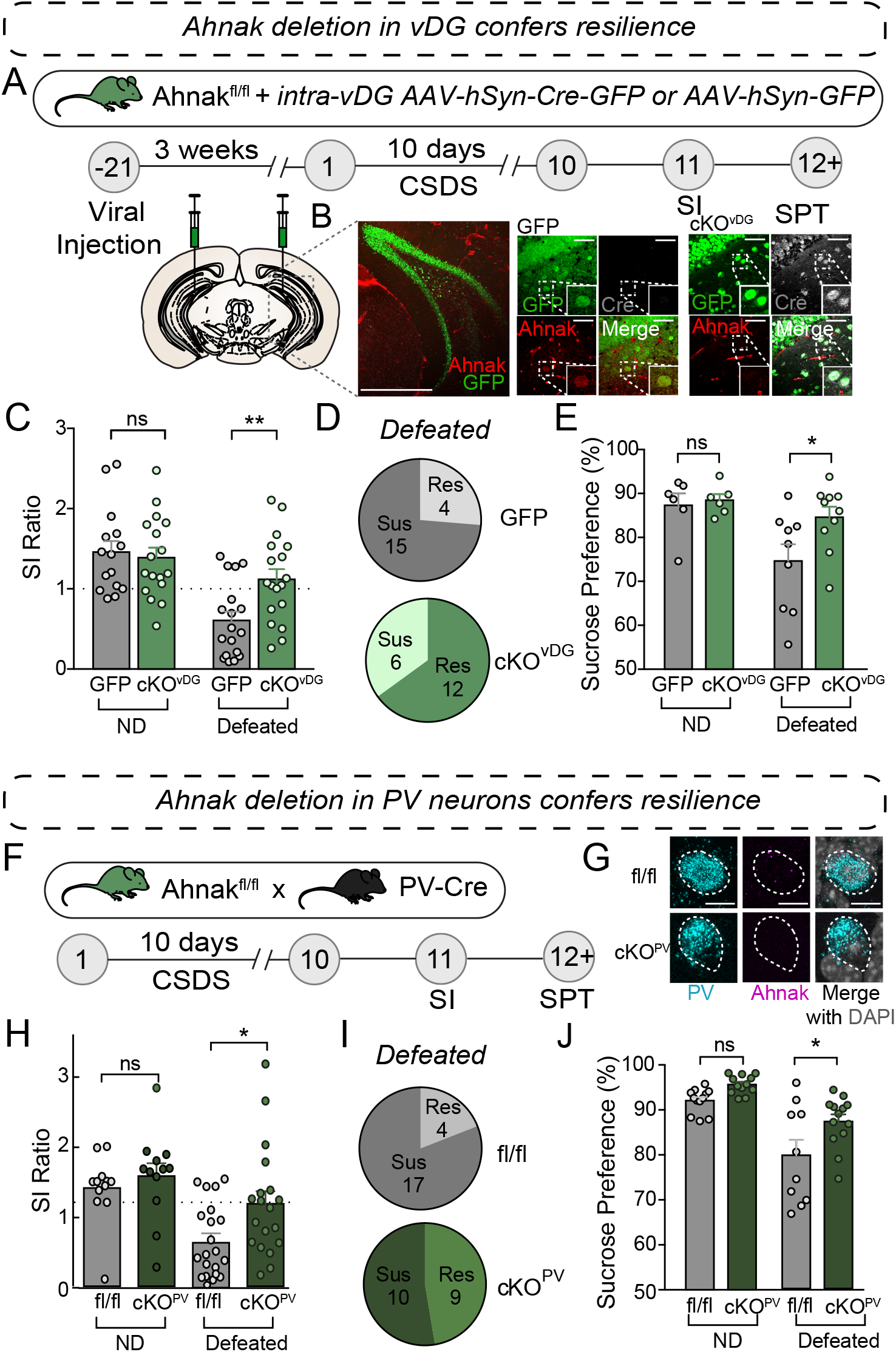
Ahnak deletion in vDG or PV neurons confers resilience to CSDS. **(A)** Diagram and timeline for conditional knockout approach using stereotaxic AAV injections of AAV-Cre-GFP or AAV-GFP control. (**B**) Representative confocal image of coronal brain section stained for GFP and Ahnak, depicting deletion of Ahnak selectively in vDG neurons. Scale Bar, 500 µm (left), 50 µm (right). **(C)** Ahnak cKO^vDG^ mice display a higher SI ratio than GFP-controls (Cont) (two-way ANOVA, Non-defeat (ND): Cont (n=16) and cKO^vDG^ (n=17), Defeated: Cont (n=19) and cKO^vDG^ (n=18)). **(D)** Pie charts showing the number of resilient and susceptible animals resulting after CSDS. **(E)** Ahnak cKO^vDG^ mice display decreased stress-induced anhedonic-like behavior in the sucrose preference test after CSDS (two-way ANOVA, ND: Cont (n=6) and cKO^vDG^ (n=6), Defeated: Cont (n=9) and cKO^vDG^ (n=10)). **(F)** Behavioral timeline and diagram of conditional knockout approach for Ahnak cKO^PV^. (**G**) Confocal images showing absence of Ahnak mRNA in PV^vDG^ neurons in mice with a conditional deletion in PV neurons (cKO^PV^). **(H)** After CSDS, cKO^PV^ mice display a higher SI ratio compared to control mice (fl/fl) (two-way ANOVA, ND: fl/fl (n=12) and cKO^PV^ (n=12), Defeated: fl/fl (n=21) and cKO^PV^ (n=19)). **(I)** Pie charts showing the number of resilient and susceptible animals resulting after CSDS. **(J)** Ahnak cKO^PV^ mice display increased sucrose preference after CSDS compared to controls (two-way ANOVA, ND: fl/fl (n=12) and cKO^PV^ (n=12), Defeated: fl/fl (n=10) and cKO^PV^ (n=13)). Data are expressed as mean ± SEM and individual data points are depicted. *Post-hoc* Bonferroni’s multiple comparisons were used for ANOVA. \**p* < 0.05, \*\**p* < 0.01 and ns, nonsignificant.

We next investigated whether Ahnak deletion selectively in PV neurons affects stress-induced behavioral responses. To delete Ahnak selectively in PV neurons, we generated offspring from crossing floxed Ahnak mice with a PV neuron-specific Cre recombinase line (cKO^PV^) (Figure 5F), which results in PV-selective Ahnak KO(20) (Figure 5G). We exposed Ahnak cKO^PV^ mice (f/f; Cre-positive) and control mice (f/f; Cre-negative) to CSDS and performed SI test and SPT (Figure 5F). Defeated Ahnak cKO^PV^ mice showed an increased SI ratio compared to the defeated control fl/fl mice (Figure 5H and Supplementary Figure 5B), yielding a greater portion of resilient mice (Figure 5I). Furthermore, defeated Ahnak cKO^PV^ mice display increased sucrose consumption compared to the defeated control group suggesting an attenuation of the CSDS-induced anhedonic-like behavior (Figure 5J). Non-defeated Ahnak cKO^PV^ mice and control mice display comparable behaviors in SI test (Figure 5H, Supplementary Figure 5B) and SPT (Figure 5J). Astoundingly, these results are consistent with those found with conditional deletion of Ahnak in the vDG (cKO^vDG^). Altogether, these data suggest that the increases in Ahnak expression observed in PV^vDG^ of susceptible mice (Figure 4E-G) is required for generating behavioral susceptibility from CSDS.

Since Ahnak and increased PV activity are both required for behavioral susceptibility, we next aimed to assess whether Ahnak deletion in PV neurons affects the activity of PV neurons, we performed whole-cell current-clamp recordings of PV^vDG^ using acute hippocampal slices from Ahnak cKO^PV^ and control mice (Supplementary Figure 6A). The firing frequency induced by current injection in PV neurons in Ahnak cKO^PV^ mice was significantly reduced compared to the control group (Supplementary Figure 6B-C), suggesting a role of Ahnak in the modulation of PV firing. Analyses of action potential properties indicate that Ahnak cKO^PV^ does not influence the voltage threshold, action potential amplitude and afterhyperpolarization (AHP), but it increases the half-amplitude width (Supplementary Figure 6D-H). These results implicate that the decrease of excitability of PV neurons by Ahnak deletion may contribute to behavioral resilience or impede susceptibility after CSDS.

## Discussion

Here, we demonstrate that PV^vDG^ neuronal activity and alterations in hippocampal Ahnak expression are critical for generating susceptibility or resilience to CSDS. Our study complements previous studies using rodent chronic stress paradigms to identify various cellular and molecular adaptations including ion channels, synaptic proteins, transcription factors, microRNAs and epigenetic regulators in key neural circuits regulating stress susceptibility or resilience including amygdalar, thalamic, and hippocampal circuits as well as catecholaminergic circuits (11, 12, 55). In this study, we identified a causal relationship of neuronal activity of PV^vDG^ in behavioral susceptibility or resilience to CSDS. We also identified that many divergent gene expression patterns emerged between resilient mice and susceptible mice as compared to non-defeated mice. For instance, DEGs associated with oxidative phosphorylation and EIF2 signaling are largely decreased in resilient mice but increased in susceptible mice compared to non-defeated mice (Figure 3K, L). However, DEGs associated with synaptogenesis or mTOR signaling comprised of a mixture of upregulated or downregulated genes across both resilient and susceptible mice (Figure 3M and Supplementary Fig 3B). These results suggest that active molecular changes involved in mitochondrial function, energy metabolism, protein translation and synaptic plasticity underlie adaptations of PV neuronal firing in stress-susceptible and resilient groups. We also identified divergent alterations of Ahnak in hippocampal lysates and PV^vDG^ (Figure 4). However, the pattern of firing changes of PV neurons is somewhat different from that of gene changes. PV neuronal firing is increased only in susceptible mice, but the firing rate in resilient mice is comparable to that of non-defeated controls (Figure 1), reminiscent of previous observations in VTA neurons (56-58). Only stress-susceptible mice display increased firing of DA neurons projecting from the VTA to the NAc and decreased firing of DA neurons projecting from the VTA to the mPFC (56, 57). Although significantly more genes are regulated in the VTA and NAc in resilient mice compared with susceptible mice (10), resilient mice display control-level firing activity in both of the aforementioned circuits (58). This adaptation of firing activity in resilient mice has been explained by homeostatic adaptation of intrinsic properties through up-regulated hyperpolarization-activated current as an excitatory driving force in conjunction with upregulation of potassium channels as an inhibitory driving force for intrinsic excitability of neurons (58). Thus, the DEGs observed in resilient mice may evoke a homeostatic control mechanism to similarly maintain the firing of PV neurons at a comparable rate with non-defeated mice.

Previously, we identified Ahnak as an endogenous regulator of L-type VGCCs in mice (20). Human genetic studies implicate altered function of L-type VGCCs in the pathophysiology of multiple psychiatric disorders including major depressive disorder, bipolar disorder, schizophrenia and autism spectrum (59-64). L-type VGCCs have been implicated in the rapid antidepressant actions of ketamine (65) and scopolamine (66). These observations raise a potential connection between Ahnak-mediated pathways and stress-induced depression-like behavior and its amelioration by antidepressants. In this study, we have observed that Ahnak level is reduced or elevated in the hippocampal PV neurons selectively in resilient or susceptible mice, respectively (Figure 4), and deletion of Ahnak in PV neurons facilitates behavioral resilience after CSDS (Figure 5). It is possible that these alterations in Ahnak contribute to, at least in part, the regulation of PV neuronal firing. This pivotal role of Ahnak in PV neurons for CSDS-induced behavior is presumably given by its regulation of L-type VGCCs. The N-terminal region of Ahnak binds to Ca_v_1.2, an L-type pore-forming α1 subunit, and its C-terminal region scaffolds the β subunit of VGCC and the p11/Anxa2 complex (20). Cell surface expression of Ca_v_1.2 and Ca_v_1.3 and L-type calcium current were significantly reduced in Ahnak KO neurons compared to wild-type controls (20). Because L-type VGCC-mediated calcium signaling mediates nuclear gene expression, synaptic plasticity and homeostatic control of neuronal circuitry (27), altered levels of Ahnak in PV neurons likely modulate calcium signaling-mediated neuronal adaptations. Notably, gene expression of most L-type VGCC subunits are not altered in comparison between stress groups with the exception of Cacnb1 gene (p=0.0244 for Res vs Sus, Supplementary Table 2). Thus, we suggest a model that Ahnak alterations potentially alter trafficking of L-type VGCCs to the plasma membrane and thereby alter channel activity, possibly resulting in gene alterations relevant to mitochondria, protein synthesis and synaptogenesis pathways.

In the brain, Ahnak scaffolds p11, alterations of which is highly implicated in the pathophysiology and antidepressant actions (67). Previously, we observed that proteins, but not mRNA levels, of p11 and Anxa2 were highly destabilized in the absence of Ahnak (20), implicating that the p11-mediated pathways may also be involved in adaptations of PV neurons in response to CSDS. In this study, we have found that p11 mRNA is also significantly increased in susceptible group compared to non-defeat control (mean RPKM 14.810 in Sus vs 11.144 in non-defeat, p= 0.0385) or resilient mice (mean RPKM 14.810 in Sus vs 10.126 in Res, p= 0.0058) in the TRAP dataset (Supplementary Table 2). P11 plays a role in gene expression by regulating SMACA3, a chromatin remodeling factor, and possibly together with Supt6, an RNA polymerase II binding partner and also known as a histone chaperone (16, 68, 69). Thus, nuclear p11 pathways may contribute to the gene alterations associated with mitochondria, protein synthesis and synaptogenesis pathways that we identified in this study. In addition to the nuclear roles, recent studies indicate roles for p11 in intrinsic membrane excitability, which is mediated by gene regulation of Kv3.1β potassium channels in hippocampal PV neurons (70) or HCN2 channels (hyperpolarization activated cyclic nucleotide gated potassium and sodium channel) in cholinergic interneurons in the nucleus accumbens (32). Importantly, Kv3.1β inhibition has been suggested to play a role in the therapeutic actions of selective serotonin reuptake inhibitor (SSRI) antidepressants by suppressing PV^vDG^ neurons via Gi-coupled signaling (70). Consistent with these observations, we have also observed suppression of PV neuron firing in resilient mice as well as ketamine-treated susceptible mice (Figure 1F). Taken together, these studies altogether suggest the PV inhibition as a common cellular mechanism associated with biological resilience as well as pharmacological actions of two different classes of antidepressants.

Interestingly, a recent study by Medrihan et al. suggested that decreased Kv3.1 channel and p11 function in PV^vDG^ would confer susceptibility after stress (71). Although Ahnak cKO^PV^ mice show resilience after CSDS (Figure 5F-J), p11 cKO^PV^ mice show stress-susceptibility after SSDS(71), likely through many other binding partners of p11 in PV^vDG^ including mGluR5, SMARCA3, Supt6 and 5-HT5A (16, 19, 70). Additionally, the timing of modulation of PV^vDG^ or vDG network activity prior, during, and after stress likely affect behavioral outcome. We show in this study that inhibition each day of chronic stress (Figure 2A-D), as well as acute inhibition after stress (Figure 2E-G), results in stress-resilience. However, Medrihan et al. observed that acute chemogenetic inhibition of vDG^PV^ during SSDS increased stress-susceptibility in the social interaction test. Further studies monitoring the activity of PV^vDG^ neurons during CSDS are necessary to fully understand their involvement in CSDS-induced behavioral responses. Importantly, most CSDS studies thus far primarily use male mice, limiting the implications arising from these studies. In recent years, however, CSDS models have been adapted for use in female mice (72-75). Because of gender differences in the prevalence of mood disorders (76, 77), the use of these newer models and testing our findings in females is warranted.

It was recently discovered that increased adult hippocampal neurogenesis inhibits a population of stress responsive GC in vDG, conferring stress resilience (47). Interestingly, Ahnak deletion promotes hippocampal neurogenesis (78). CSDS-induced hippocampal Ahnak reductions might promote hippocampal neurogenesis as an alternative mechanism for stress resilience. Notably, ∼25% reduction of Ahnak protein in hippocampal lysates in resilient mice (Figure 4C) cannot be explained solely by Ahnak reduction in PV interneurons, which constitutes at best ∼ 5% of total hippocampal neurons (79). Thus, our findings do not eliminate a potential involvement of Ahnak expressed in other neuronal types or non-neuronal cells in stress resilience. In fact, Ahnak is highly expressed in endothelial cells in blood vessels in the hippocampus (20, 80), and tight junctions therein provide the property of the blood-brain barrier (81). Intriguingly, endothelial cells have been implicated as a target of CSDS, and CSDS-induced dysfunction of the BBB has been suggested as a mechanism underlying stress susceptibility (45). Thus, further understanding of the function of Ahnak in other cell types will advance our understanding of stress resilience. Altogether, our study establishes a foundation supporting Ahnak and PV interneurons as potential targets for manipulation of stress-related hippocampal physiology or neuropsychiatric disorders such as MDD.

## Acknowledgements

We greatly appreciate Dr. Paul Greengard for his generous support for our research program. We thank Claire Song for technical assistance and Elizabeth Griggs for graphic presentation. We also thank Rada Norinsky and The Rockefeller University Transgenic and Reproductive Technology Center for their excellent *in vitro* fertilization and embryo transfer services. This work was supported by the National Institute of Mental Health of the National Institutes of Health under Award Number R01MH121763 (to Y.K.), the Black Family foundation, and the Fisher Center for Alzheimer’s Research Foundation.

## Author contributions

D.L.B. and Y.K. conceptualized and designed the entire study. L.M. performed and analyzed *in vivo* electrophysiological experiments. L.M. and K.M. performed and analyzed *ex vivo* electrophysiological experiments. D.L.B. and M.X.C. performed and analyzed all behavioral experiments. D.L.B. performed all stereotaxic surgeries. D.L.B., M.X.C., and J.J. performed and analyzed all immunoblotting experiments. D.L.B. and M.X.C. performed and analyzed all RNAscope experiments. M.X.C., J.L., and E.P.A. performed tissue collection for TRAP. M.X.C., J.L., and W.W. performed analysis for TRAP experiments. D.L.B. and Y.K. wrote the manuscript and designed figures with collective input from all authors. Y.K. supervised the entire study.

## Data and materials availability

RNA-seq data from Figure 3 have been deposited and are available from GEO (accession number: GSE184027). Additional data related to this paper may be requested from the authors.

## Figure legends

**Supplementary Figure 1. Interaction time in non-defeated control, resilient and susceptible groups after CSDS**. Time spent in the interaction zone without (-) and with (+) aggressor (two-way repeated measures ANOVA, Non-defeated (ND) (n=23), Resilient (Res) (n=23) and Susceptible (Sus) (n=15)).

**Supplementary Figure 2. Chemogenetic modulation of PV^vDG^ predispose divergent behavioral outcome in response to social defeat stress. (A, B) Gi DREADD experiments. (A)** Repeated CNO-treatment during CSDS resulted in increased time in the interaction zone with the aggressor compared to time in the empty interaction zone in hM4Di mice, while no difference was seen in mCherry controls (two-way repeated measures ANOVA, Non-defeat: mCherry (n=8) and hM4Di (n=12), Defeated: mCherry (n=15) and hM4Di (n=14)). **(B)** Acute CNO-treatment in susceptible mice resulted in increased time in the interaction zone with aggressor in hM4Di mice compared to mCherry controls and to treatment-free SI-tests (SI-1 and SI-3) (two-way repeated measures ANOVA, mCherry (n=6) and h4MDi (n=9)). **(C-H)** Gq DREADD experiments. **(C, D)** Acute activation during SSDS does not alter SI ratio (C) or time spent in the interaction zone (D) in hM3Dq or mCherry control mice (two-way repeated measures ANOVA, Non-defeat: mCherry (n=8) and h3MDq (n=10), Defeated: mCherry (n=13) and h3MDq (n=18)). **(E)** A trend of decreased time spent in the interaction zone in hM3Dq mice compared to mCherry controls was found after repeated injections of CNO post-SSDS (two-way repeated measures ANOVA, mCherry (n=12) and hM3Dq (n=15)). (**F, G**) Non-SSDS mice (social defeat-free) had no alterations in SI ratio (F) or time spent in the interaction zone (G) before or after repeated injections of CNO (two-way repeated measures ANOVA, mCherry (n=8) and hM3Dq (n=10)). (**H**) Repeated injections of CNO did not alter sucrose preference in non-SSDS mice (unpaired two-tailed t-test, mCherry (n=8) and hM3Dq (n=9)). Data are expressed as mean ± SEM, and individual data points are depicted. *Post-hoc* Bonferroni’s multiple comparisons were used for ANOVA. \**p*<0.05, \*\**p*<0.01.

**Supplementary Figure 3. Molecules associated with mitochondrial dysfunction and mTOR signaling are altered in hippocampal PV neurons in divergently behaving mouse groups after CSDS. (A, B)** Heat maps of DEGs involved in mitochondrial dysfunction and mTOR signaling. Mean value of gene expression was used (Non-defeated (n=4), Res (n=4) and Sus (n=3)).

**Supplementary Figure 4. Chronic stress induces alterations of Ahnak expression in the hippocampus. (A)** Hippocampal Ahnak is inversely correlated with time spent in interaction zone with aggressor mouse (pearson *r* : *r* = -0.5071, *p<*0.0001, n=61 mice) (**B**) Ahnak mRNA RNAScope puncta is inversely correlated with time in interaction zone with aggressor (pearson *r* : *r* = -0.5071, *p<*0.0001, n=10 mice).

**Supplementary Figure 5. Ahnak deletion in vDG or PV neurons confers resilience to CSDS**. (**A**) In non-defeated conditions, both control and Ahnak cKO^vDG^ groups display higher amount of time spent in the interaction zone with an aggressor compared to the time without aggressor. After CSDS, control mice display significantly lower amount of time spent in the interaction zone with an aggressor compared to the time without aggressor, but Ahnak cKO^vDG^ mice display equal amounts of time in the interaction zone with or without an aggressor (two-way repeated measures ANOVA, Non-defeat: Control (n=15) and cKO^vDG^ (n=17), Defeated: Control (n=18) and KO^vDG^ (n=18)). (**B)** CSDS induces decreased interaction time during the aggressor session compared to empty-cage session in control group (floxed Ahnak mice, fl/fl), while the effect of CSDS on interaction time with an aggressor is abolished in Ahnak cKO^PV^ mice (two-way repeated measures ANOVA, Non-defeat: fl/fl (n=12) and cKO^PV^ (n=12), Defeated: fl/fl (n=21) and cKO^PV^ (n=19)). Data are expressed as mean ± SEM, and individual data points are depicted. *Post-hoc* Bonferroni’s multiple comparisons were used for ANOVA. \**p*<0.05, \*\*\**p*<0.001, \*\*\*\**p*<0.0001 and ns, nonsignificant.

**Supplementary Figure 6. The effect of PV neuron-selective Ahnak deletion on physiological properties of PV neurons in the vDG. (A)** Schematic of whole-cell patch clamp. (**B**) Representative traces from whole-cell current-clamped PV neurons in the vDG of fl/fl and Ahnak cKO^PV^ mice showing the action potential (AP) firing of the cells in response to a 500 pA step of injected current. **(C)** AP frequency of PV neurons in the vDG is reduced in Ahnak cKO^PV^ mice at incremental steps of injected current (two-way ANOVA, fl/fl (n=10 neurons/4 mice) and cKO^PV^ (n=9 neurons/4 mice)). Data are expressed as mean ± SEM. *Post-hoc* Bonferroni’s multiple comparisons were used for ANOVA. **(D)** Representative single APs in PV^vDG^ neurons from control (fl/fl, black) and Ahnak cKO^PV^ (green) mice. **(E-H)** AP properties of PV^vDG^ neurons in Ahnak cKO^PV^ and control mice. Ahnak cKO^PV^ does not influence the voltage threshold (**E**), AP amplitude (**F**), Afterhyperpolarization (**H**) but increases the half-amplitude width (**G**). Two-tailed unpaired *t*-test, 8 neurons/4 mice for control (fl/fl) and 9 neurons/mice for cKO^PV^ for each experiment). Data are expressed as mean ± SEM, and individual data points are depicted. \**p*<0.05, \*\**p*<0.01 and ns, nonsignificant.

## References

1. Hollon NG, Burgeno LM, Phillips PE (2015): Stress effects on the neural substrates of motivated behavior. Nature neuroscience. 18:1405–1412.

2. Nestler EJ, Barrot M, DiLeone RJ, Eisch AJ, Gold SJ, Monteggia LM (2002): Neurobiology of depression. Neuron. 34:13–25.

3. Kendler KS, Karkowski LM, Prescott CA (1999): Causal relationship between stressful life events and the onset of major depression. American journal of psychiatry. 156:837–841.

4. Kessler RC (1997): The effects of stressful life events on depression. Annu Rev Psychol. 48:191–214.

5. Syed SA, Nemeroff CB (2017): Early Life Stress, Mood, and Anxiety Disorders. Chronic Stress (Thousand Oaks). 1:1–16.

6. McEwen BS (2012): Brain on stress: how the social environment gets under the skin. Proc Natl Acad Sci U S A. 109 Suppl 2:17180–17185.

7. Southwick SM, Vythilingam M, Charney DS (2005): The psychobiology of depression and resilience to stress: implications for prevention and treatment. Annu Rev Clin Psychol. 1:255–291.

8. Cryan JF, Mombereau C (2004): In search of a depressed mouse: utility of models for studying depression-related behavior in genetically modified mice. Mol Psychiatry. 9:326–357.

9. Nasca C, Bigio B, Zelli D, Nicoletti F, McEwen BS (2015): Mind the gap: glucocorticoids modulate hippocampal glutamate tone underlying individual differences in stress susceptibility. Mol Psychiatry. 20:755–763.

10. Krishnan V, Han MH, Graham DL, Berton O, Renthal W, Russo SJ, et al. (2007): Molecular adaptations underlying susceptibility and resistance to social defeat in brain reward regions. Cell. 131:391–404.

11. Han MH, Nestler EJ (2017): Neural Substrates of Depression and Resilience. Neurotherapeutics. 14:677–686.

12. Cathomas F, Murrough JW, Nestler EJ, Han MH, Russo SJ (2019): Neurobiology of Resilience: Interface Between Mind and Body. Biol Psychiatry. 86:410–420.

13. Campbell S, Macqueen G (2004): The role of the hippocampus in the pathophysiology of major depression. Journal of psychiatry & neuroscience : JPN. 29:417–426.

14. Fanselow MS, Dong HW (2010): Are the dorsal and ventral hippocampus functionally distinct structures? Neuron. 65:7–19.

15. Malberg JE, Eisch AJ, Nestler EJ, Duman RS (2000): Chronic antidepressant treatment increases neurogenesis in adult rat hippocampus. The Journal of neuroscience : the official journal of the Society for Neuroscience. 20:9104–9110.

16. Oh YS, Gao P, Lee KW, Ceglia I, Seo JS, Zhang X, et al. (2013): SMARCA3, a chromatin-remodeling factor, is required for p11-dependent antidepressant action. Cell. 152:831–843.

17. Umschweif G, Greengard P, Sagi Y (2021): The dentate gyrus in depression. Eur J Neurosci. 53:39–64.

18. Shuto T, Kuroiwa M, Sotogaku N, Kawahara Y, Oh YS, Jang JH, et al. (2020): Obligatory roles of dopamine D1 receptors in the dentate gyrus in antidepressant actions of a selective serotonin reuptake inhibitor, fluoxetine. Mol Psychiatry. 25:1229–1244.

19. Lee KW, Westin L, Kim J, Chang JC, Oh YS, Amreen B, et al. (2015): Alteration by p11 of mGluR5 localization regulates depression-like behaviors. Mol Psychiatry. 20:1546–1556.

20. Jin J, Bhatti DL, Lee KW, Medrihan L, Cheng J, Wei J, et al. (2020): Ahnak scaffolds p11/Anxa2 complex and L-type voltage-gated calcium channel and modulates depressive behavior. Mol Psychiatry. 25:1035–1049.

21. Haase H, Alvarez J, Petzhold D, Doller A, Behlke J, Erdmann J, et al. (2005): Ahnak is critical for cardiac Ca(V)1.2 calcium channel function and its beta-adrenergic regulation. FASEB J. 19:1969–1977.

22. Matza D, Badou A, Kobayashi KS, Goldsmith-Pestana K, Masuda Y, Komuro A, et al. (2008): A scaffold protein, AHNAK1, is required for calcium signaling during T cell activation. Immunity. 28:64–74.

23. Hotka M, Cagalinec M, Hilber K, Hool L, Boehm S, Kubista H (2020): L-type Ca(2+) channel-mediated Ca(2+) influx adjusts neuronal mitochondrial function to physiological and pathophysiological conditions. Sci Signal. 13.

24. Guzman JN, Ilijic E, Yang B, Sanchez-Padilla J, Wokosin D, Galtieri D, et al. (2018): Systemic isradipine treatment diminishes calcium-dependent mitochondrial oxidant stress. J Clin Invest. 128:2266–2280.

25. D’Arco M, Dolphin AC (2012): L-type calcium channels: on the fast track to nuclear signaling. Sci Signal. 5:pe34.

26. Dolmetsch RE, Pajvani U, Fife K, Spotts JM, Greenberg ME (2001): Signaling to the nucleus by an L-type calcium channel-calmodulin complex through the MAP kinase pathway. Science. 294:333–339.

27. Simms BA, Zamponi GW (2014): Neuronal voltage-gated calcium channels: structure, function, and dysfunction. Neuron. 82:24–45.

28. Autry AE, Adachi M, Nosyreva E, Na ES, Los MF, Cheng PF, et al. (2011): NMDA receptor blockade at rest triggers rapid behavioural antidepressant responses. Nature. 475:91–95.

29. Ma XC, Dang YH, Jia M, Ma R, Wang F, Wu J, et al. (2013): Long-lasting antidepressant action of ketamine, but not glycogen synthase kinase-3 inhibitor SB216763, in the chronic mild stress model of mice. PLoS One. 8:e56053.

30. Kim JW, Autry AE, Na ES, Adachi M, Bjorkholm C, Kavalali ET, et al. (2021): Sustained effects of rapidly acting antidepressants require BDNF-dependent MeCP2 phosphorylation. Nature neuroscience. 24:1100–1109.

31. Golden SA, Covington HE, 3rd, Berton O, Russo SJ (2011): A standardized protocol for repeated social defeat stress in mice. Nat Protoc. 6:1183–1191.

32. Cheng J, Umschweif G, Leung J, Sagi Y, Greengard P (2019): HCN2 Channels in Cholinergic Interneurons of Nucleus Accumbens Shell Regulate Depressive Behaviors. Neuron. 101:662–672 e665.

33. Medrihan L, Sagi Y, Inde Z, Krupa O, Daniels C, Peyrache A, et al. (2017): Initiation of Behavioral Response to Antidepressants by Cholecystokinin Neurons of the Dentate Gyrus. Neuron. 95:564–576 e564.

34. Stark E, Eichler R, Roux L, Fujisawa S, Rotstein HG, Buzsaki G (2013): Inhibition-induced theta resonance in cortical circuits. Neuron. 80:1263–1276.

35. Senzai Y, Buzsaki G (2017): Physiological Properties and Behavioral Correlates of Hippocampal Granule Cells and Mossy Cells. Neuron. 93:691–704 e695.

36. Toader O, Forte N, Orlando M, Ferrea E, Raimondi A, Baldelli P, et al. (2013): Dentate gyrus network dysfunctions precede the symptomatic phase in a genetic mouse model of seizures. Front Cell Neurosci. 7:138.

37. Heiman M, Kulicke R, Fenster RJ, Greengard P, Heintz N (2014): Cell type-specific mRNA purification by translating ribosome affinity purification (TRAP). Nat Protoc. 9:1282–1291.

38. Roussarie JP, Yao V, Rodriguez-Rodriguez P, Oughtred R, Rust J, Plautz Z, et al. (2020): Selective Neuronal Vulnerability in Alzheimer’s Disease: A Network-Based Analysis. Neuron. 107:821–835 e812.

39. Chen S, Zhou Y, Chen Y, Gu J (2018): fastp: an ultra-fast all-in-one FASTQ preprocessor. Bioinformatics. 34:i884–i890.

40. Dobin A, Davis CA, Schlesinger F, Drenkow J, Zaleski C, Jha S, et al. (2013): STAR: ultrafast universal RNA-seq aligner. Bioinformatics. 29:15–21.

41. McCarthy DJ, Chen Y, Smyth GK (2012): Differential expression analysis of multifactor RNA-Seq experiments with respect to biological variation. Nucleic Acids Res. 40:4288–4297.

42. Love MI, Huber W, Anders S (2014): Moderated estimation of fold change and dispersion for RNA-seq data with DESeq2. Genome Biol. 15:550.

43. Schmittgen TD, Livak KJ (2008): Analyzing real-time PCR data by the comparative C(T) method. Nat Protoc. 3:1101–1108.

44. Berton O, McClung CA, Dileone RJ, Krishnan V, Renthal W, Russo SJ, et al. (2006): Essential role of BDNF in the mesolimbic dopamine pathway in social defeat stress. Science. 311:864–868.

45. Menard C, Pfau ML, Hodes GE, Kana V, Wang VX, Bouchard S, et al. (2017): Social stress induces neurovascular pathology promoting depression. Nature neuroscience. 20:1752–1760.

46. Kheirbek MA, Drew LJ, Burghardt NS, Costantini DO, Tannenholz L, Ahmari SE, et al. (2013): Differential control of learning and anxiety along the dorsoventral axis of the dentate gyrus. Neuron. 77:955–968.

47. Anacker C, Luna VM, Stevens GS, Millette A, Shores R, Jimenez JC, et al. (2018): Hippocampal neurogenesis confers stress resilience by inhibiting the ventral dentate gyrus. Nature. 559:98–102.

48. Scharfman HE, Myers CE (2012): Hilar mossy cells of the dentate gyrus: a historical perspective. Front Neural Circuits. 6:106.

49. Kim J, Farchione T, Potter A, Chen Q, Temple R (2019): Esketamine for Treatment-Resistant Depression - First FDA-Approved Antidepressant in a New Class. New England journal of medicine. 381:1–4.

50. Kraushaar U, Jonas P (2000): Efficacy and stability of quantal GABA release at a hippocampal interneuron-principal neuron synapse. The Journal of neuroscience : the official journal of the Society for Neuroscience. 20:5594–5607.

51. Struber M, Jonas P, Bartos M (2015): Strength and duration of perisomatic GABAergic inhibition depend on distance between synaptically connected cells. Proceedings of the National Academy of Sciences of the United States of America. 112:1220–1225.

52. Roth BL (2016): DREADDs for Neuroscientists. Neuron. 89:683–694.

53. Heiman M, Schaefer A, Gong S, Peterson JD, Day M, Ramsey KE, et al. (2008): A translational profiling approach for the molecular characterization of CNS cell types. Cell. 135:738–748.

54. Wang F, Flanagan J, Su N, Wang LC, Bui S, Nielson A, et al. (2012): RNAscope: a novel in situ RNA analysis platform for formalin-fixed, paraffin-embedded tissues. J Mol Diagn. 14:22–29.

55. Russo SJ, Murrough JW, Han MH, Charney DS, Nestler EJ (2012): Neurobiology of resilience. Nature neuroscience. 15:1475–1484.

56. Cao JL, Covington HE, 3rd, Friedman AK, Wilkinson MB, Walsh JJ, Cooper DC, et al. (2010): Mesolimbic dopamine neurons in the brain reward circuit mediate susceptibility to social defeat and antidepressant action. Journal of Neuroscience. 30:16453–16458.

57. Chaudhury D, Walsh JJ, Friedman AK, Juarez B, Ku SM, Koo JW, et al. (2013): Rapid regulation of depression-related behaviours by control of midbrain dopamine neurons. Nature. 493:532–536.

58. Friedman AK, Walsh JJ, Juarez B, Ku SM, Chaudhury D, Wang J, et al. (2014): Enhancing depression mechanisms in midbrain dopamine neurons achieves homeostatic resilience. Science. 344:313–319.

59. Green EK, Grozeva D, Jones I, Jones L, Kirov G, Caesar S, et al. (2010): The bipolar disorder risk allele at CACNA1C also confers risk of recurrent major depression and of schizophrenia. Mol Psychiatry. 15:1016–1022.

60. (2013): Identification of risk loci with shared effects on five major psychiatric disorders: a genome-wide analysis. Lancet. 381:1371–1379.

61. Bhat S, Dao DT, Terrillion CE, Arad M, Smith RJ, Soldatov NM, et al. (2012): CACNA1C (Cav1.2) in the pathophysiology of psychiatric disease. Progress in neurobiology. 99:1–14.

62. Pinggera A, Lieb A, Benedetti B, Lampert M, Monteleone S, Liedl KR, et al. (2015): CACNA1D de novo mutations in autism spectrum disorders activate Cav1.3 L-type calcium channels. Biological psychiatry. 77:816–822.

63. Schizophrenia Working Group of the Psychiatric Genomics C (2014): Biological insights from 108 schizophrenia-associated genetic loci. Nature. 511:421–427.

64. Liu Y, Blackwood DH, Caesar S, de Geus EJ, Farmer A, Ferreira MA, et al. (2011): Meta-analysis of genome-wide association data of bipolar disorder and major depressive disorder. Mol Psychiatry. 16:2–4.

65. Lepack AE, Bang E, Lee B, Dwyer JM, Duman RS (2016): Fast-acting antidepressants rapidly stimulate ERK signaling and BDNF release in primary neuronal cultures. Neuropharmacology. 111:242–252.

66. Yu H, Li M, Shen X, Lv D, Sun X, Wang J, et al. (2018): The Requirement of L-Type Voltage-Dependent Calcium Channel (L-VDCC) in the Rapid-Acting Antidepressant-Like Effects of Scopolamine in Mice. The international journal of neuropsychopharmacology. 21:175–186.

67. Svenningsson P, Kim Y, Warner-Schmidt J, Oh YS, Greengard P (2013): p11 and its role in depression and therapeutic responses to antidepressants. Nat Rev Neurosci. 14:673–680.

68. Lu H, Xie Y, Tran L, Lan J, Yang Y, Murugan NL, et al. (2020): Chemotherapy-induced S100A10 recruits KDM6A to facilitate OCT4-mediated breast cancer stemness. J Clin Invest. 130:4607–4623.

69. Umschweif G, Medrihan L, McCabe KA, Sagi Y, Greengard P (2021): Activation of the p11/SMARCA3/Neurensin-2 pathway in parvalbumin interneurons mediates the response to chronic antidepressants. Mol Psychiatry. 26:3350–3362.

70. Sagi Y, Medrihan L, George K, Barney M, McCabe KA, Greengard P (2020): Emergence of 5-HT5A signaling in parvalbumin neurons mediates delayed antidepressant action. Mol Psychiatry. 25:1191–1201.

71. Medrihan L, Umschweif G, Sinha A, Reed S, Lee J, Gindinova K, et al. (2020): Reduced Kv3.1 Activity in Dentate Gyrus Parvalbumin Cells Induces Vulnerability to Depression. Biological psychiatry. 88:405–414.

72. Yohn CN, Dieterich A, Bazer AS, Maita I, Giedraitis M, Samuels BA (2019): Chronic non-discriminatory social defeat is an effective chronic stress paradigm for both male and female mice. Neuropsychopharmacology. 44:2220–2229.

73. Takahashi A, Chung JR, Zhang S, Zhang H, Grossman Y, Aleyasin H, et al. (2017): Establishment of a repeated social defeat stress model in female mice. Sci Rep. 7:12838.

74. Harris AZ, Atsak P, Bretton ZH, Holt ES, Alam R, Morton MP, et al. (2018): A Novel Method for Chronic Social Defeat Stress in Female Mice. Neuropsychopharmacology. 43:1276–1283.

75. Newman EL, Covington HE, 3rd, Suh J, Bicakci MB, Ressler KJ, DeBold JF, et al. (2019): Fighting Females: Neural and Behavioral Consequences of Social Defeat Stress in Female Mice. Biological psychiatry. 86:657–668.

76. Riecher-Rossler A (2017): Sex and gender differences in mental disorders. Lancet Psychiatry. 4:8–9.

77. Seney ML, Sibille E (2014): Sex differences in mood disorders: perspectives from humans and rodent models. Biol Sex Differ. 5:17.

78. Shin JH, Kim YN, Kim IY, Choi DH, Yi SS, Seong JK (2015): Increased Cell Proliferations and Neurogenesis in the Hippocampal Dentate Gyrus of Ahnak Deficient Mice. Neurochemical research. 40:1457–1462.

79. Pelkey KA, Chittajallu R, Craig MT, Tricoire L, Wester JC, McBain CJ (2017): Hippocampal GABAergic Inhibitory Interneurons. Physiological reviews. 97:1619–1747.

80. Gentil BJ, Benaud C, Delphin C, Remy C, Berezowski V, Cecchelli R, et al. (2005): Specific AHNAK expression in brain endothelial cells with barrier properties. Journal of cellular physiology. 203:362–371.

81. Daneman R, Prat A (2015): The blood-brain barrier. Cold Spring Harb Perspect Biol. 7:a020412.

