## Supplementary figures and images for "Molecular and cellular adaptations in hippocampal parvalbumin neurons mediate behavioral responses to chronic social stress"

### Supplementary Figure 1

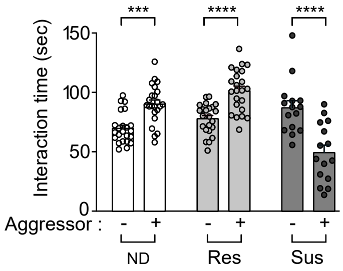


**Supplementary Figure 1**

### Supplementary Figure 2

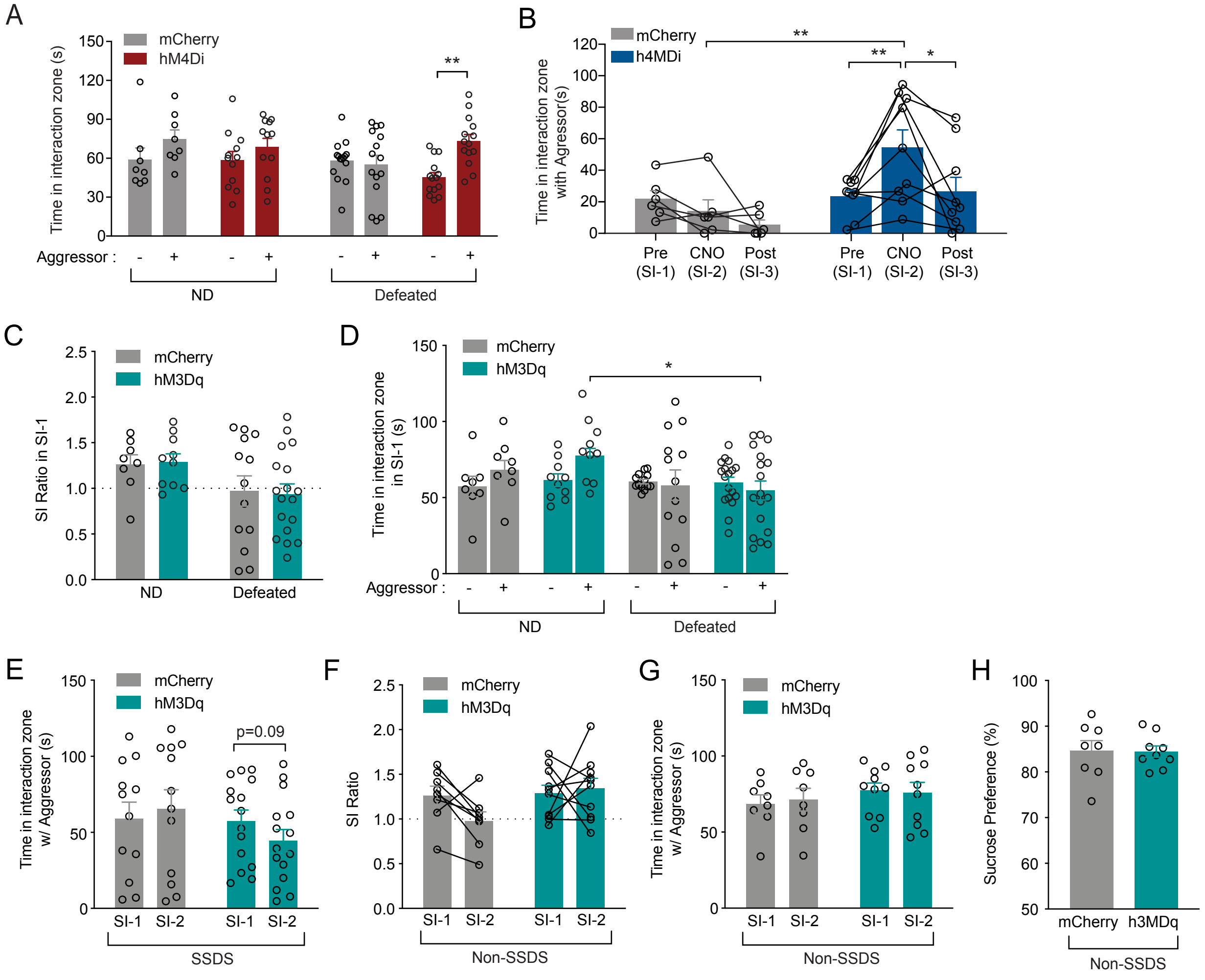


**Supplementary Figure 2**

### Supplementary Figure 3

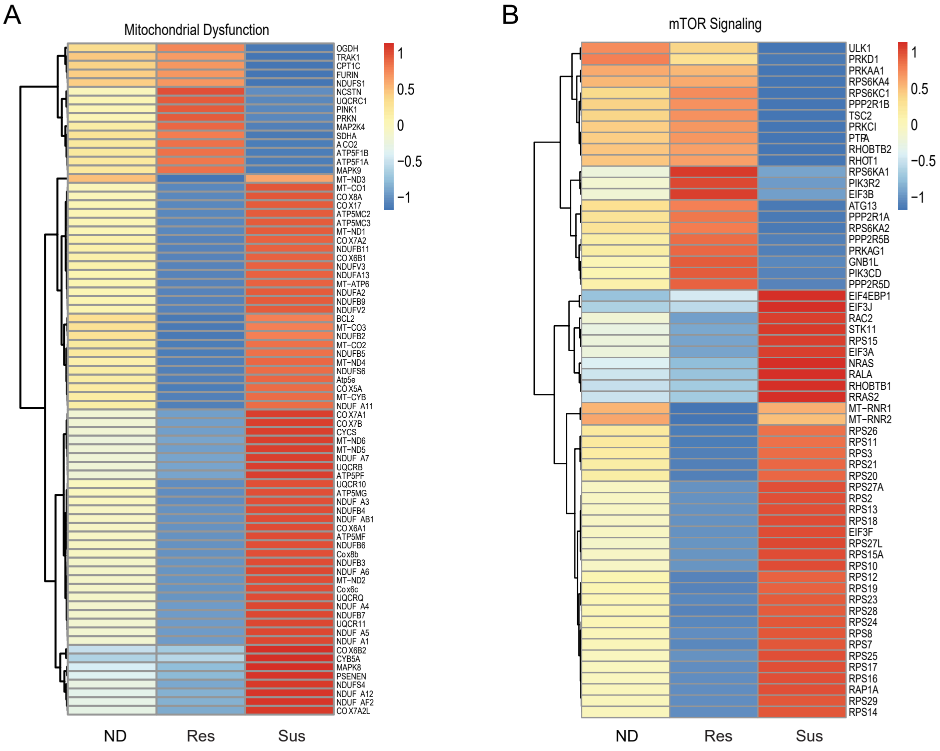


**Supplementary Figure 3**

### Supplementary Figure 4

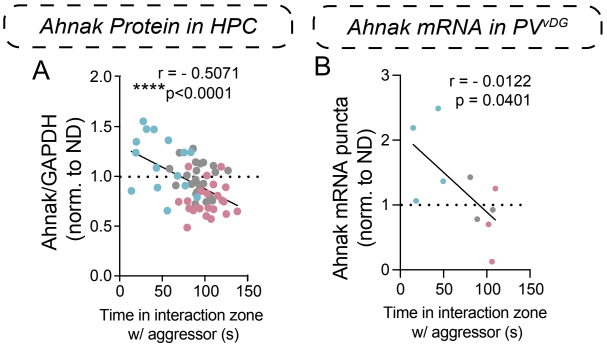


**Supplementary Figure 4**

### Supplementary Figure 5

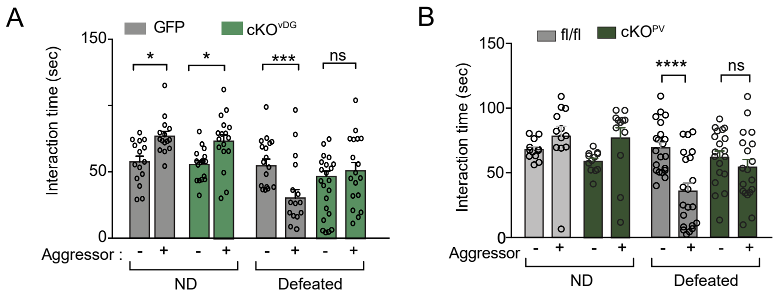


**Supplementary Figure 5**

### Supplementary Figure 6

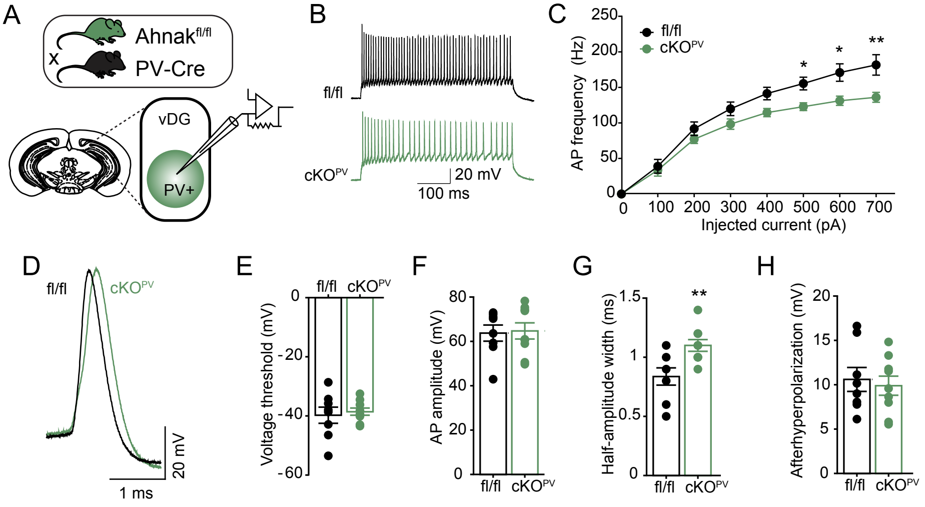


**Supplementary Figure 6**
