## Supplementary Table 1 for "Molecular and cellular adaptations in hippocampal parvalbumin neurons mediate behavioral responses to chronic social stress"

**Supplementary Table 1** Summary of statistical analysis (related to all Figures).

| Figure | | Experimental variables | Statistical test | Results |
|---|---|---|---|---|
| 1 | **B** | group (Non-defeat, Res, Sus) | one-way ANOVA | group: F(2,58)=40.82, *p*<0.0001 |
| | **F** | group (Cont, Res, Sus, Sus+Ket) | one-way ANOVA | group: F (3, 138) = 7.671, *p* < 0.0001 |
| | **G** | group (Cont, Res, Sus, Sus+Ket) | one-way ANOVA | group: F (3, 199) = 6.597, *p* = 0.0003 |
| | **H** | group (Cont, Res, Sus, Sus+Ket) | one-way ANOVA | group: F (3, 131) = 8.773, *p* < 0.0001 |
| | **I** | group (Cont, Res, Sus, Sus+Ket) | one-way ANOVA | group: F (3, 290) = 3.611, *p* = 0.0138 |
| 2 | **C** | group (mCherry, hM4Di); treatment (non-defeat, defeated) | two-way ANOVA | group: F(1,45)=3.693, *p*=0.0610; treatment: F(1,45)=0.04982, *p*=0.8244; interaction, F(1,45)=10.23, *p*=0.0025 |
| | **F** | group (mCherry, hM4Di); treatment (non-defeat, defeated) | two-way ANOVA | group: F(1,48)=6.808, *p*=0.0121; treatment: F(1,48)=3.207, *p*=0.0796; interaction: F(1,48)=4.616, *p*=0.0368 |
| | **H** | time (Pre, CNO, Post); group (mCherry, hM4Di) | two-way repeated measures ANOVA | time: F(2,26) = 1.805, *p*=0.1845; group: F(1,13)=2.834, *p*=0.1161; interaction: F(2,26)=4.945, *p*=0.0151; subject: F(13,26)=1.935, *p*=0.0737 |
| | **I** | hM4Di vs. mCherry | Two-tailed unpaired student's *t*-test | t(14)=3.174, *p*=0.0068 |
| | **M** | time (SI-1, SI-2); group (mCherry, hM3Dq) | two-way repeated measures ANOVA | time: F(1,25)=2.348, *p*=0.1380; group: F(1,25)=0.6389, *p*=0.4316; interaction: F(1,25)=2.756, *p*=0.1094; subject: F(25,25)=3.687, *p*=0.0009 |
| | **O** | hM3Dq vs. mCherry | two-tailed unpaired student's *t*-test | t(25)=2.147, *p*=0.0416 |

| | | | | |
|---|---|---|---|---|
| 4 | **C** | group (Non-defeat, Res, Sus) | one-way ANOVA | group: F(2,59)=21.48, *p*<0.0001 |
| | **D** | SI Ratio vs. Ahnak protein | Pearson's Correlation | R2=0.1471, *p*=0.0008 |
| | **F** | group (Non-defeat, Res, Sus) | Per Cell: Krusal-Wallis test, Dunn's multiple comparison | Per Cell:<br>Krusal-Wallis statistic: 38.89, *p*<0.0001; ND vs. Res: *p*=0.001; ND vs Sus: *p*=0.0092 |
| | **G** | SI Ratio vs. normalized Ahnak puncta per cell | Pearson's Correlation | R2=0.4911, *p*=0.0240 |
| 5 | **C** | group (fl/fl, $cKO^{vDG}$); treatment (non-defeat, defeated) | two-way ANOVA | group: F(1,64)=20.48, *p*<0.0001; treatment: F(1,64)=3.190, *p*=0.0788; interaction, F(1,64)=5.543 *p*=0.0216 |
| | **E** | group (fl/fl, $cKO^{vDG}$); treatment (non-defeat, defeated) | two-way ANOVA | group: F(1,27)=7.450, *p*<0.0110; treatment: F(1,27)=3.377, *p*=0.0771, interaction, F(1,27)=2.126, *p*=0.1564 |
| | **H** | group (fl/fl, $cKO^{PV}$); treatment (non-defeat, defeated) | two-way ANOVA | group: F(1,60)=4.796, *p*=0.0110; treatment: F(1,60)=12.92, *p*=0.0007; interaction, F(1,60)=1.289, *p*=0.2608 |
| | **J** | group (fl/fl, $cKO^{PV}$); treatment (non-defeat, defeated) | two-way ANOVA | group: F(1,43)=9.779, *p*=0.0032; treatment: F(1,43)=34.76, *p*<0.0001; interaction, F(1,43)=1.239, *p*=0.2719 |
| S1 | | time (absence, presence of aggressor); group (non-defeat, Res, Sus) | two-way repeated measures ANOVA | time: F(1,58) = 0.5697, *p*=0.4534; group: F(2,58) = 11.10, *p*<0.0001; interaction: F(2,58)=42.24, *p*<0.0001; subject: F(58,58)=1.590, *p*=0.0401 |
| S2 | **A** | time (absence, presence of aggressor); group (non-defeat mCherry, non-defeat hM4Di, | two-way repeated measures ANOVA | time: F(1,45)=9.803, *p*=0.0031; group: F(3,45)=0.9129, *p*=0.4423; interaction, F(3,45)=3.042, *p*=0.0384; subject: F(45,45)=1.274, *p*=0.2101. |

| | | | | |
|---|---|---|---|---|
| | | defeated mCherry, defeated hM4Di) | | |
| | **B** | time (SI-1, SI-2, SI-3); group (mCherry, hM4Di) | two-way repeated measures ANOVA | time: F(2,26)=4.214, *p*=0.0260; group: F(1,13)=5.703, *p*=0.0328; interaction, F(2,26)=4.945, *p*=0.0151; subject: F(31,31)=4.620, *p*<0.0001 |
| | **C** | group (mCherry, hM3Dq); treatment (non-defeat, defeated); | two-way ANOVA | group: F(1,45)=0.0002492, *p*=0.9875; treatment: F(1,45)=5.623, *p*=0.0221; interaction, F(1,45)=0.0474, *p*=0.8287 |
| | **D** | time (aggressor, no agressor); group (non-defeat mCherry, non-defeat hM43q, defeated mCherry, defeated hM3Dq); | two-way repeated measures ANOVA | time: F(1,45)=1.465, *p*=0.2324; group: F(3,45)=1.310, *p*=0.2830; interaction, F(3,45)=1.752, *p*=0.1700; subject, F(45,45)=1.445, *p*=0.1105 |
| | **E** | Time (SI-1, SI-2); group (mCherry, hM3Dq) | two-way repeated measures ANOVA | time: F(1,25)=0.4731, *p*=0.4979, group: F(1,25)=0.8439, *p*=0.3671; interaction: F(1,25)=4.408, *p*=0.0460, subject: F(25,25)=6.997, *p*<0.0001 |
| | **F** | Time (SI-1, SI-2); group (mCherry, hM3Dq) | two-way repeated measures ANOVA | time: F(1,16)=1.172, *p*=0.2951; group: F(1,16)=3.803; *p*=0.0689; interaction, F(1,16) = 2.534, *p* = 0.1310; F(16,16)=0.88997, *p*=0.5824 |
| | **G** | Time (SI-1, SI-2); group (mCherry, hM3Dq) | two-way repeated measures ANOVA | time: F(1,16)=0.008782, *p*=0.9265, group: F(1,16)=1.111, *p*=0.3074; interaction: F(1,16)=0.1347, *p*=0.7184; subject: F(16,16)=0.9884, *p*=0.5092 |
| | **H** | hM3Dq vs. mCherry | two-tailed unpaired student's *t*-test | t(15)=0.09129, *p*=0.9285 |
| S4 | **A** | Time in interaction zone w/ Agg. Ahnak protein | Pearson's Correlation | R2=0.2572, *p*<0.0001 |
| | **B** | Time in Agg Zone vs. normalized Ahnak puncta per cell | Pearson's Correlation | R2=0.4282 *p*=0.0401 |

| | | | | |
|---|---|---|---|---|
| S5 | **A** | time (aggressor, no agressor); group (non-defeat fl/fl, non-defeat cKO-vDG, defeated defeated cKO-vDG); | two-way repeated measures ANOVA | time: $F(1,64)=0.2.057$, $p=0.1564$; group: $F(3,64)= 9.806$, $p<0.0001$; interaction, $F(3,64)=11.19$, $p<0.0001$; subject, $F(64,64)=1.566$, $p=0.0375$ |
| | **B** | time (aggressor, no agressor); group (non-defeat fl/fl, non-defeat cKO-PV, defeated defeated cKO-PV); | two-way repeated measures ANOVA | time: $F(1,60)=0.8963$, $p=0.3476$; group: $F(3,60)= 4.853$, $p=0.0043$; interaction, $F(3,60)=9.608$, $p<0.0001$; subject, $F(60,60)=1.291$, $p=0.1625$ |
| S6 | **C** | group (fl/fl, $cKO^{PV}$); injected current | two-way ANOVA | firing at the same injected current : $F (7, 122) = 97.53$, $P < 0.0001$; firing at increased injected current : $F (1, 122) = 31.68$, $P < 0.0001$; Interaction: $F (7, 122) = 1.791$, $P = 0.0949$ |
| | **E** | group (fl/fl, $cKO^{PV}$) | unpaired two-tailed student's *t* test | $p = 0.6777$, $t(15)=0.4238$ |
| | **F** | group (fl/fl, $cKO^{PV}$) | unpaired two-tailed student's *t* test | $p = 0.8552$, $t(15)=0.1856$ |
| | **G** | group (fl/fl, $cKO^{PV}$) | unpaired two-tailed student's *t* test | $p = 0.0085$, $t(15)=3.024$ |
| | **H** | group (fl/fl, $cKO^{PV}$) | unpaired two-tailed student's *t* test | $p = 0.6841$, $t(15)=0.4148$ |
